# PEDOT:PSS Microparticles for Extrudable and Bioencapsulating Conducting Granular Hydrogel Bioelectronics

**DOI:** 10.1101/2025.05.25.655149

**Authors:** Anna P. Goestenkors, Justin S. Yu, Jae Park, Yuqing Wu, Cielo J. Vargas Espinoza, Lianna C. Friedman, Somtochukwu S. Okafor, Tianran Liu, Suman Chatterjee, Avishek Debnath, Barbara A. Semar, Cayleigh P. O’Hare, Riley M. Alvarez, Srikanth Singamaneni, Baranidharan Raman, Alexandra L. Rutz

## Abstract

Conducting hydrogels are promising materials for forming physiomimetic bioelectronic interfaces to monitor and stimulate biological activity. However, most developed materials are non-microporous and possess fixed shapes, both of which can limit the integration of cells and tissues with devices. In non-conducting biomaterials, materials fabrication strategies imparting microporosity and dynamic mechanical properties have been shown to support cell infiltration and support biointerfaces of various geometries. Specifically, granular hydrogels have enabled encapsulating, conformal, and injectable interfaces through these features. However, granular hydrogels remain largely unexplored as conducting biomaterials. We present methods for fabricating spherical, poly(3,4-ethylenedioxythiophene):poly(styrene sulfonate) (PEDOT:PSS) hydrogel microparticles. When densely packed, these microparticles form a conducting granular hydrogel with microporosity as well as shear-thinning and self-healing dynamic mechanical properties. The PEDOT:PSS granular hydrogel can be extruded and maintain structure post-3D printing. Modulating microparticle PSS content achieves high granular hydrogel conductivity (137 S/m), and microparticles exhibit excellent cytocompatibility (>98% viability). Finally, we demonstrate utility as bioencapsulating electrodes for electrophysiological monitoring. These results highlight the functionality of our PEDOT:PSS conducting granular hydrogel, suggesting its potential as 3D printed bioencapsulating electrodes, 3D tissue engineering scaffolds for monitoring encapsulated cells, and injectable therapies for enhanced cell recruitment and tissue regeneration combined with electronic stimulation.

## Introduction

Bioelectronics are used as a continuous, label-free method to monitor biological activities of cells and tissues. Examples include wearable devices for monitoring muscle loading and fatigue, implantable devices for monitoring of cardiac activity and detection of arrhythmias, and in vitro devices for detection of action potentials produced from neurons for better understanding of cellular functions^1–3^. However, traditional bioelectronic devices rely on metals, silicon, plastics, and glass as the direct material interface with biology. Typical devices are characterized by stiffnesses exceeding that of native tissue as well as discrete and planar biointerfaces^4^. These device attributes lead to limitations such as inability to generate electronics-integrated yet physiomimetic cell culture models in vitro for studying disease or screening drug compounds^5^. In vivo, limited tissue integration with the device may lead to an exacerbated foreign body response and low signal-to-noise ratios limiting the quality and duration of biological activity recordings^6^.

Hydrogels are being investigated as replacements to these constituent materials for bioelectronic devices with tissue-mimetic properties to improve the biointerface. Such hydrogels can include those solely selected to support surrounding biology or those that also serve as functional parts of the device itself. Functional hydrogels include those that are electronically conducting and facilitate ion-to-electron transduction when used as electrodes for electrophysiological monitoring or stimulation for guiding cell differentiation, behavior, and activity^7–9^. While findings on hydrogel-based bioelectronics have been highly promising, a majority of these reports focus on hydrogels that are non-microporous (herein referred to as ‘bulk hydrogels’) and/or have static networks resulting in hydrogels of fixed shapes^4^. Microporosity in hydrogels provides a means by which cells can integrate within the material, for either 3D cell culture in vitro or host cell infiltration in vivo. Static shape refers to a structure which cannot undertake alternative form factors without permanent material failure. Alternatively, dynamic mechanical properties, including the ability to flow upon shear and self-heal once applied forces are removed, have been shown to be useful for adapting to a variety of biointerfaces through conformability (close contact with surface of tissue), injectability (enabling minimally invasive implantation), and encapsulation (enveloping cells or tissues).

Granular hydrogels have become a widely used technique in non-conducting biomaterials to provide both microporosity as well as dynamic mechanical properties in hydrogels. Granular hydrogels are composed of densely packed micron-scale hydrogel particles (i.e. microparticles) (Figure 1a). In this densely packed state, or what is commonly referred to as the jammed state, the material possesses an interconnected microporous structure facilitated by the void spaces left between densely packed microparticles. This interconnected microporous structure facilitates cell encapsulation, infiltration, migration, spreading, and proliferation in both in vitro and in vivo applications^10,11^. Additionally, interactions between the microparticles result in shear-thinning and self-healing dynamic mechanical properties^12,13^. These dynamic mechanical properties allow for the granular hydrogel to be extruded and maintain the resultant structure, be injected into cavities, and conform to topographically diverse surfaces^10,14^. Exploiting both of these properties, researchers have applied these materials as injectable biomaterial scaffolds for in situ tissue regeneration^15,16^, 3D printed implantable structures for in vivo drug release^17^, and 3D bioprinted cell-laden in vitro cell culture scaffolds^13,18,19^.

**Figure 1.**
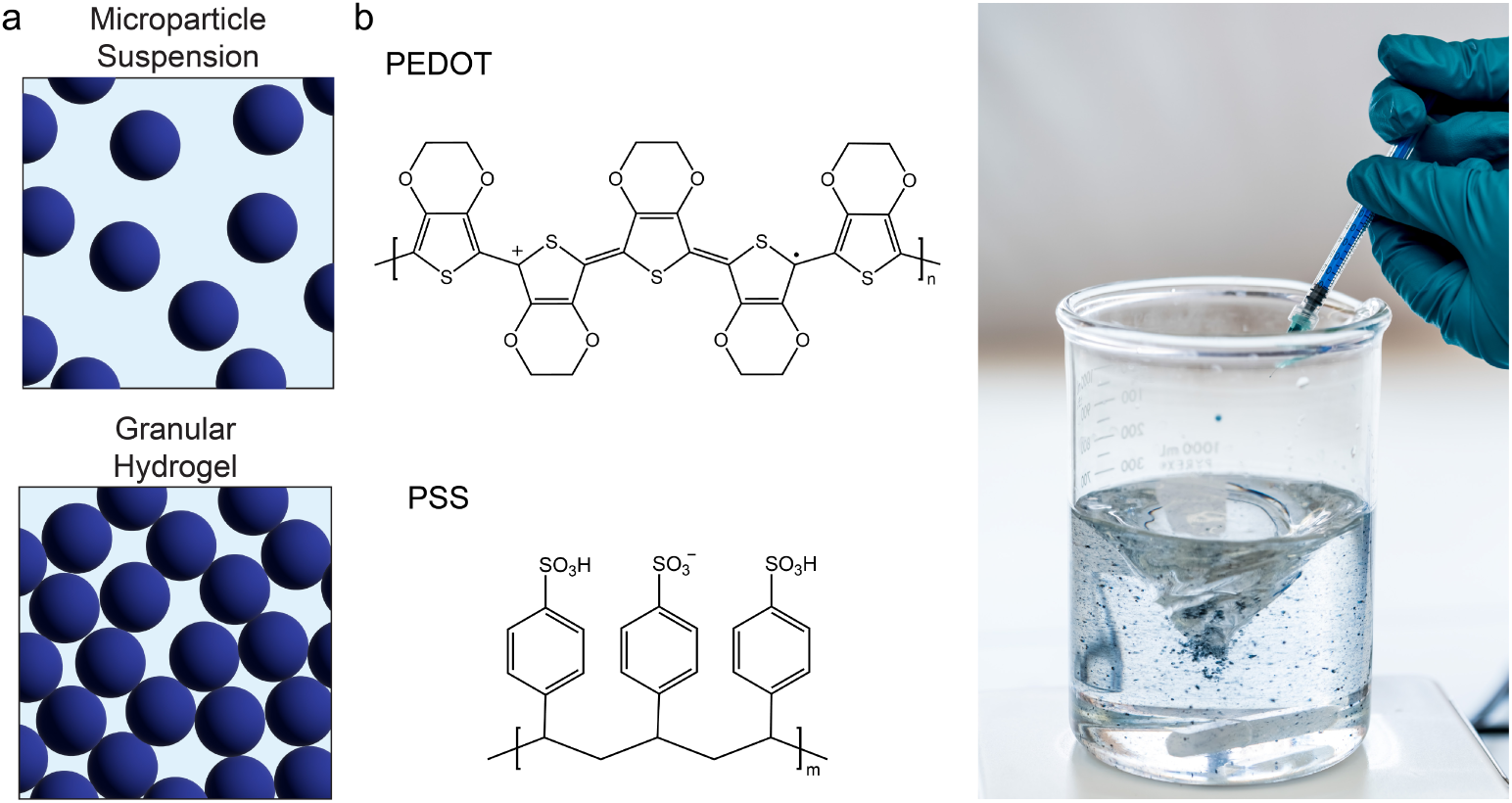
PEDOT:PSS microparticles fabricated by batch water-in-oil emulsion are proposed for generation of conducting granular hydrogels. A) Granular hydrogels are comprised of densely packed hydrogel microparticles. After microparticle suspensions in water are densely packed, the material possesses an interconnected microporous structure as well as shear-thinning and self-healing dynamic mechanical properties. B) The conducting polymer PEDOT:PSS is used to generate microparticles. An aqueous colloidal dispersion of PEDOT:PSS is sheared into microdroplets when injected into a spinning oil phase.

While granular hydrogels have been increasingly investigated in non-conducting materials, there are only a few studies to date on electronically conducting granular hydrogels^20–25^. Bringing this form factor to conducting materials would expand material capabilities and possible bioapplications, as highlighted in the works above. In order to fabricate granular hydrogels, two criteria must be fulfilled: (1) hydrogel microparticles are generated with diameters of tens to hundreds of micrometers and (2) particles are packed to achieve the jammed state^26^. While not mandated requirements for granular hydrogel formation, particle shape and size homogeneity can benefit applications of both extrudability and cell-material interactions. When compared to heterogeneous, angular microparticles, spherical microparticles required lower extrusion forces which corresponded to higher cell viability of encapsulated cells when the material and cells were extruded simultaneously^27^. Additionally, spherical microparticles resulted in smaller, cell-size porosity when compared to that of heterogeneous, angular microparticles^27^. An additional criterion to consider for conducting granular hydrogels specifically is the resulting electronic conductivity. While a desired value is application-dependent, materials with conductivities generally exceeding 1 S/m have been successfully demonstrated as functional components of bioelectronic devices^28^.

The first step towards the generation of conducting granular hydrogels is the fabrication of the building block itself, conducting hydrogel microparticles. Candidate materials include hydrogels comprised of polymers that are intrinsically conducting (doped conjugated polymers) or non-conducting hydrogels loaded with a percolating network of nano-sized conducting particles. Common techniques employed for microparticle fabrication include microfluidic emulsions, batch emulsions, electrospraying, mechanical fragmentation, or lithography techniques^26^. Among prior works of conducting granular hydrogels specifically, mechanical fragmentation, microfluidic emulsions, and electrospraying have been presented thus far. Microfluidic emulsions and electrospraying yielded uniformly shaped particles and monodisperse size. However, reported strategies have shown lower conductivities (<1 S/m) likely due to reliance on a non-conducting hydrogel carrier loaded with metallic or conducting polymer nanoparticles^20,21^. Additionally, higher concentrations of conducting polymer added during the fabrication process were shown to impact microparticle shape and size, ultimately leading to larger, more fibrous structures rather than spheres^20^. Mechanical fragmentation has been performed on bulk, millimeter-scale hydrogels comprised solely of conducting polymer. Higher conductivities were achieved in this granular hydrogel (∼10 S/m), yet generated particles have irregular shapes and heterogeneous sizes.^22,25^

In this manuscript, we aimed to expand investigations of these limited number of works on conducting granular hydrogels. We selected pure conducting polymer as the material choice given its promise for higher electronic conductivities. While microfluidic emulsions yield uniform particle populations, generation of microfluidic devices and keeping channels from clogging can limit microparticle production rates. Instead, we sought a simpler fabrication method of a batch emulsion where an aqueous phase containing polymer is dispersed in an oil phase under high shear followed by the introduction of a crosslinking trigger ^27,29–31^. Microparticle fabrication methods were developed for the conducting polymer poly(3,4-ethylenedioxythiophene):poly(styrene sulfonate) (PEDOT:PSS) (Figure 1b) followed by evaluations of how particles can successfully generate granular hydrogels and exhibit properties desirable for bioelectronic applications including extrudability, cytocompatibility, tissue-conformability, and use as part of an electrode device.

## Results and Discussion

### PEDOT:PSS Microparticles Fabricated Via Batch Emulsion Using Heated Oil Phase

Of available conducting polymers, we selected PEDOT:PSS for our investigations. The conducting polymer PEDOT:PSS is known for its excellent aqueous stability and conductivity as thin films^32^ and has been used to fabricate injectable^33^, cast^34^, and 3D printed conducting hydrogels^35^. PEDOT:PSS has also been used to form nanometer-scale particles^36–38^ and less commonly micrometer-scale particles comprised solely of conducting polymer^39,40^. The former falls outside our desired size range and the latter reports have not been investigated for generating granular hydrogels. As previously stated, we aimed to fabricate hydrogel microparticles using a simple batch water-in-oil emulsion which involves dispersing an aqueous phase with polymer into an oil phase. PEDOT:PSS is commercially available as a colloidal aqueous dispersion containing nano-scale aggregates of this polymer complex (1.0-1.3 wt%)^41,42^. Therefore, we proposed this PEDOT:PSS colloidal aqueous dispersion could serve as the aqueous phase in the emulsion.

Synthesis of hydrogel microparticles via batch water-in-oil emulsion involves (1) shearing an aqueous polymer phase into microdroplets in an oil, (2) presenting a condition that induces crosslinking to facilitate gel formation, and (3) subsequent isolation of solid microparticles upon removal of the oil phase. There are several ways to induce crosslinking and generate solid PEDOT:PSS materials from the colloidal aqueous dispersion. These methods generally work by forming PEDOT-rich regions abundant with pi-pi stacking through concentrating the PEDOT:PSS nano-aggregates (such as through removal of its solvent or inducing phase separation) and/or removing excess PSS after applying some conditions that weaken or break down electrostatic interactions between PEDOT and PSS^39,43–45^. We sought a simple and easily applied trigger for crosslinking. Others have shown that heat treatment of the PEDOT:PSS colloidal aqueous dispersion with temperatures between 140 °C and 200 °C can lead to the formation of aqueous stable PEDOT:PSS thin films through what is likely induced phase separation resulting in PEDOT-rich insoluble regions^43^. In our own prior work, we have shown that incubation of the PEDOT:PSS colloidal dispersion at 60 °C overnight induces gelation for generation of millimeter-sized bulk gels, albeit the gels were weakly cross-linked as evident by aqueous instablility^34^. Thus, in this work, we speculated that by elevating the temperature of the oil phase under shear and subsequently adding PEDOT:PSS colloidal dispersion, we could achieve a water-in-oil emulsion in which PEDOT:PSS would undergo gelation to form aqueous stable microparticles.

Here, we investigated three temperatures – room temperature, 60 °C, and 90 °C, with the two latter temperatures motivated because of evidence of solidification in prior works. In water-in-oil emulsions, surfactants are used to stabilize microdroplets of the dispersed aqueous phase. In this work, we included 0.05 wt% of the non-ionic surfactant Span-80® in our oil phase, as it is commonly used in emulsions to improve emulsion stability^31,46^. Span-80® has a closed-cup flashpoint at temperatures greater than ∼110 °C. Therefore, we did not investigate temperatures beyond 90 °C for safety concerns. To investigate the effect of oil phase temperature on microparticle generation, we first examined for possible PEDOT:PSS gelation at these temperatures using oscillatory rheology (Figure 2, Supplementary Figure 1). As expected, neither gelation (G’ storage modulus-G” loss modulus cross-over, tan(δ) < 1) nor an increase in G’ were observed over the studied period of 60 minutes at 25 °C. At 60 °C, gelation appeared to occur near 60 minutes after G’ increased nearly 2 orders of magnitude and tan(δ) approached 1 (9.34 to 1.12). At 90 °C, fast gelation of 2-3 minutes was observed, and gelation stabilized around 20-30 minutes as evidenced by a plateau in G’ and tan(δ).

**Figure 2.**
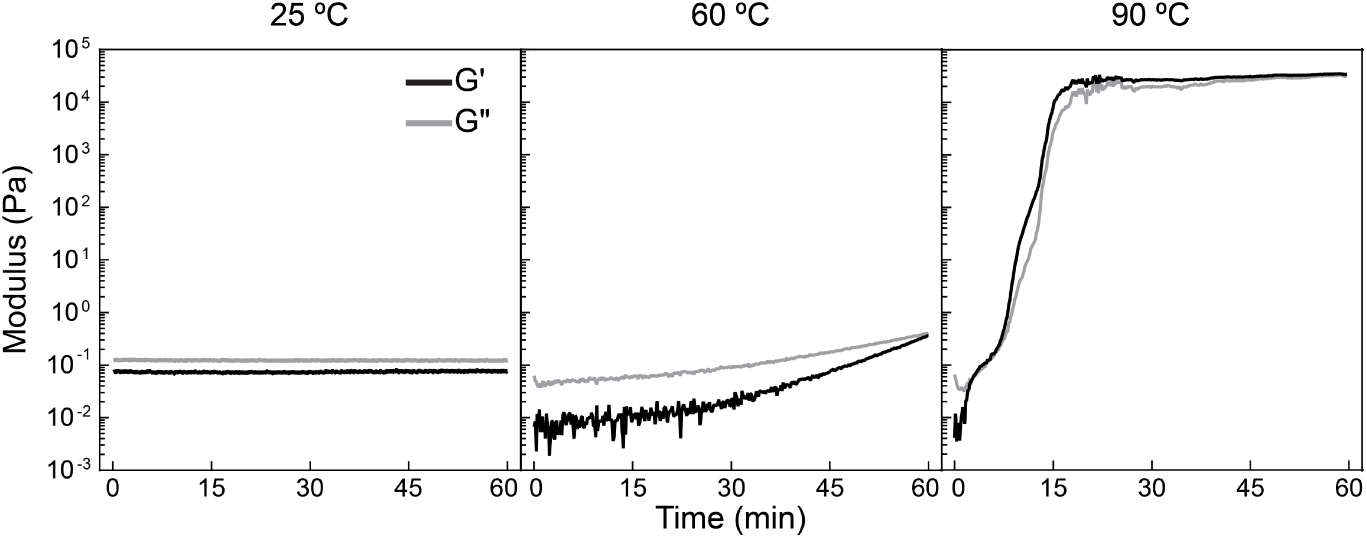
Elevated temperatures induce gelation of PEDOT:PSS colloidal dispersion with highest temperature providing fastest gelation kinetics. Oscillatory rheology of PEDOT:PSS colloidal dispersion over 60 minutes at the designated temperatures measures storage modulus (G’) and loss modulus (G”) (1% strain, 1 rad s^-1^). No gelation (G’-G” cross-over) was observed at 25 °C, nor changes in G’ and G”. Gelation occurred slowly at 60 °C with the gel point around 60 minutes (tan(δ) of 1.123). The liquid-to-gel transition (G’-G” cross-over) occurred at 90 °C within 2-3 minutes. Repeated runs are shown in Supplementary Figure 1.

To confirm heat facilitates microparticle gelation, emulsions were then conducted with the oil phase held at the various temperatures. Upon injection of the PEDOT:PSS colloidal dispersion into the spinning oil phase, generated PEDOT:PSS microdroplets were inspected to evaluate their characteristics (Figure 3a). At 25 °C, droplets were of micro-size (69 µm ± 53 µm) at 2 minutes. Over the studied period of 60 minutes, the microdroplets decreased in size by 32.66% but remained similar in color (Figure 3a, b). When attempting to remove from the oil phase by adding copious water, the PEDOT:PSS droplets diffused into the surrounding aqueous phase thus confirming unsuccessful crosslinking (Supplementary Figure 2). While a decrease in droplet size can be evidence of crosslinking, this would not be expected to be the case here for 25 °C due to failure to form aqueous stable particles; instead, it has been shown that ongoing exposure to shear and turbulence within an emulsion can decrease liquid droplet diameters^47^. Droplets from emulsions generated at 60 °C and 90 °C decreased in size over 60 minutes but appeared to darken in color (indicating a denser material) (Figure 3a,b). When isolation from the oil phase was attempted, both conditions resulted in solid particles that were aqueous stable, confirming that the addition of heat induces PEDOT:PSS gelation (Figure 4a).

**Figure 3.**
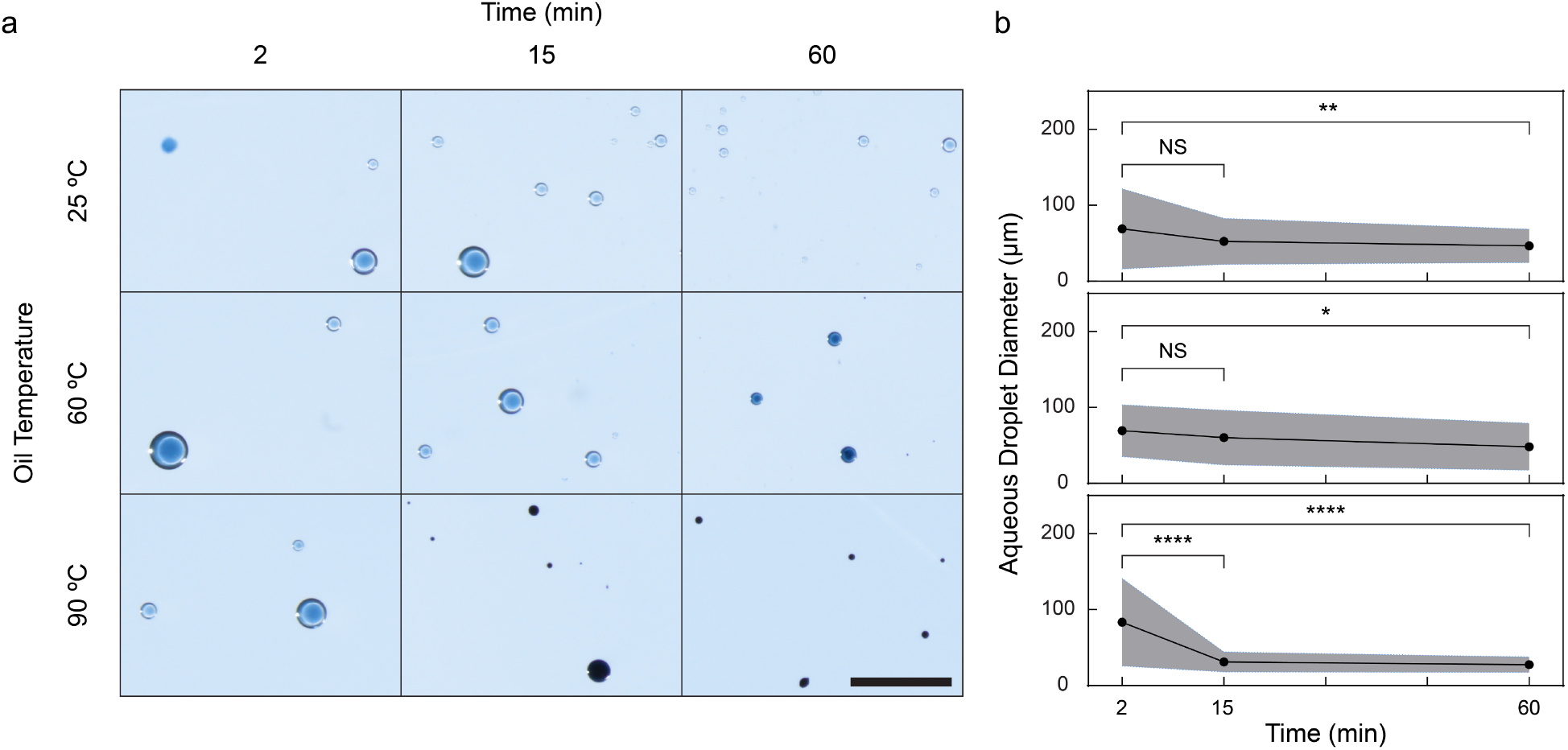
Changes in PEDOT:PSS droplets in water-in-oil emulsion indicate crosslinking at elevated temperatures. A) When PEDOT:PSS colloidal dispersion was injected into the oil phase, shear and turbulence provided by a stir bar resulted in the formation of micrometer-scale PEDOT:PSS droplets. Under continuous stirring, PEDOT:PSS droplets appeared to darken in color for 60 °C and 90 °C over 60 minutes but remained similar in color for 25 °C. Scale bar 1 mm. B) Droplet diameters decreased in all conditions, but most significantly at 90 °C. Temperatures of 25 °C, 60 °C, and 90 °C from top to bottom. N=50 droplets from 2 emulsions per condition, 25 droplets measured at each timepoint per emulsion. Plot lines denote mean and shaded areas denote standard deviation. One-way analysis of variance (ANOVA) and Tukey’s multiple comparison test. ****P≤ 0.0001 ***P ≤ 0.001 **P ≤ 0.01 *P ≤ 0.05 non-significant (NS) P > 0.05.

**Figure 4.**
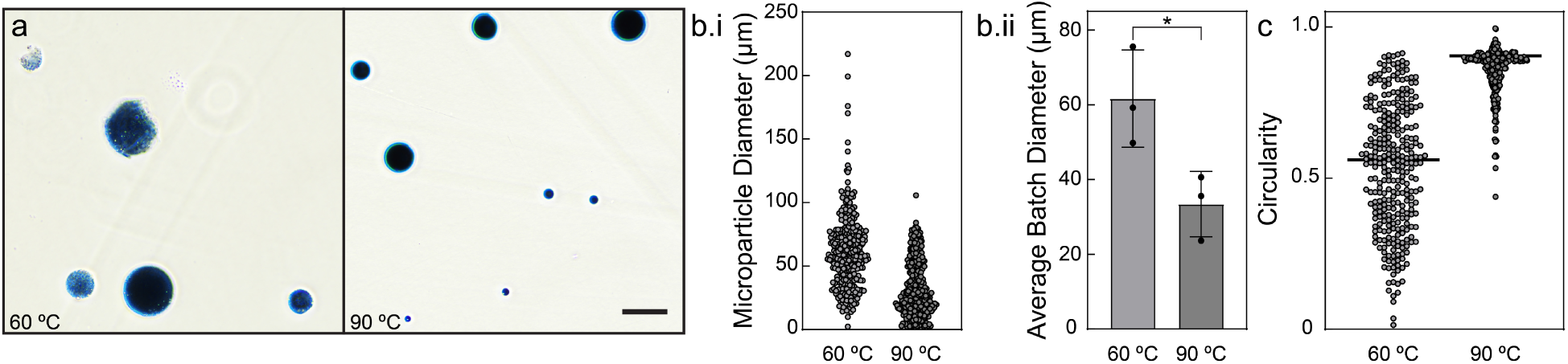
90 °C oil phase temperature results in more uniformly sized and shaped PEDOT:PSS microparticles relative to 60 °C. A) Droplets from emulsions performed at different temperatures were subjected to removal from oil phase and washing with PBS to evaluate for aqueous stability. Aqueous stable microparticles were isolated from conditions of 60 °C and 90 °C, but not 25 °C (Supplementary Figure 2). Particles in PBS from 90 °C appeared uniform in color with smooth edges while those from 60 °C were heterogeneous in color and appeared more irregular in their borders, indicating gelation may have been inconsistent. Representative images, scale bar 100 µm. Bi) Diameters of microparticle population (N=300 from 3 fabrication batches, 100 microparticles quantified per batch). Bii) Mean diameter per fabrication batch (N=3). 90 °C generated microparticles were significantly smaller than those of 60 °C (mean batch diameters of 33 µm and 62 µm, respectively). Fabrication batch values obtained by averaging values of 100 analyzed microparticles. Standard deviation presented. Unpaired t-test. ****P ≤ 0.0001 ***P ≤ 0.001 **P ≤ 0.01 *P ≤ 0.05 non-significant (NS) P > 0.05. C) Circularity values of microparticle population (N=300 from 3 fabrication batches, 100 microparticles quantified per batch). 90 °C generated microparticles had higher mean circularity than those of 60 °C (0.8575 and 0.5376, respectively, as designated by line).

Even though particles were successfully generated and isolated from emulsions at both 60 °C and 90 °C, particles from 90 °C were noticeably different by several metrics. Firstly, generated microdroplets in the emulsion appeared to become darker blue over 60 minutes and decreased in size more significantly (67.12% and 30.42% decrease in diameter for 90 °C and 60 °C, respectively; Figure 3a,b). When isolated from the oil phase and suspended in water, particles from 90 °C were about half the size as those generated from 60 °C (average batch diameters 33 µm ± 9 µm and 62 µm ± 13 µm, respectively; Figure 4b.ii). Additionally, a more uniformly sized population was observed as evident by the range of individual microparticle diameters, which was smaller for 90 °C than 60 °C (2-106 µm and 2-217 µm, respectively; Figure 4b.i). We would like to note particle size analysis relied upon optical microscopy as well as use of the image processing technique of thresholding, therefore we did not expect to detect particles less than about 1 µm. This is not of concern to the study since microparticles below this size are not relevant to the formation of granular hydrogels (require microparticles >10 µm). Isolated particles from 90 °C were uniform in color darkness and exhibited spherical shape with smooth edges resulting in a mean circularity of 0.8575 (Figure 4a,c, Supplementary Figure 3). Alternatively, particles from 60 °C appeared to have a range of color darkness with some particles even appearing translucent (Figure 4a, Supplementary Figure 4). In addition, while some microparticles exhibited spherical shapes, many irregularly shaped and rough particles were also detected and these typically exhibited circularity values <0.70 (Supplementary Figure 4b). These irregularly shaped and rough particles contributed to a lower mean circularity of 0.5376 for the particle population (Figure 4c, Supplementary Figure 4).

In all, faster and more complete gelation from 90 °C is likely contributing to these results of smaller and more uniformly shaped microparticle populations. At the time of particle isolation (60 minutes), gelation has stabilized for 90 °C whereas gelation has just started for 60 °C. Evolving gelation could explain the variance in apparent particle density (as visualized by microparticle translucence)^48^. Additionally, we have previously shown that denser crosslinking results in decreased hydrogel size^34^. It is likely that with additional time, microparticle gelation would become more complete at 60 °C and result in further decrease in microparticle diameters. In summary, due to the production of smaller microparticle populations with a more uniform shape and smaller size range, 90 °C oil temperature was defined as the optimal temperature for producing microparticles throughout the rest of this study.

### PEDOT:PSS Granular Hydrogel Exhibits Microporosity and Dynamic Mechanical Properties

The form factor of granular hydrogels arises when paths of contacting microparticles are achieved by packing particles closely together (Figure 1a). These “paths” of particles are referred to as load-bearing force chains that give rise to the material’s unique mechanical properties^26^. At the microparticle scale (>10 µm in diameter), gravitational forces become more dominant than thermal forces of the system and help maintain the densely packed state of the microparticles. Once densely packed, frictional forces at the particle-particle junctions become the most influential forces within the material^26,49^. These frictional forces and their reversible disruption and reformation give rise to the shear-thinning (decreasing viscosity in response to increased shear rate) and self-healing properties (recovers mechanical properties following removal of high strains that caused material deformation)^13,26,49^. These dynamic mechanical properties in turn give rise to the granular hydrogel’s extrudability (shear-thinning) and maintenance of extruded structures (self-healing) utilized for different applications including 3D printed tissue engineering scaffolds, 3D bioprinted cell-laden structures, and injectable materials for regenerative therapies^12,50,51^. An additional attribute of these densely packed particles is an interconnected microscale porosity beneficial for promoting cellular integration within the material. When injected into tissue voids, granular hydrogels have been shown to promote cellular infiltration, proliferation, and subsequent tissue regeneration in these wounds at a greater rate than bulk hydrogel injectables with nano-scale porosity that rely on material degradation for appreciable cell infiltration^15^. Researchers have also utilized the microporosity to encapsulate cells within the granular hydrogel pores, both as an in vitro 3D cell culture material and injectable cell therapies^16,50,52,53^.

To investigate if our PEDOT:PSS microparticles could be used to assemble a granular hydrogel, we concentrated microparticle suspensions using vacuum filtration. The microparticles formed a paste-like material that could be easily manipulated with a spatula (Figure 5a). Porosity was next evaluated by adding a fluorescent polysaccharide (fluorescein isothiocyanate–dextran, FITC-dextran) to the water prior to vacuum filtration to aid in visualization of the space among microparticles. Epifluorescence and confocal microscopy revealed microparticles surrounded by fluorescent media denoting the void spaces left between the densely packed microparticles (Figure 5b). The fluorescence also shows that these void spaces, or pores, are linked to one another, forming pathways throughout the material. These porous pathways support the formation of an interconnected microporosity typically possessed by granular hydrogels. We quantified pore sizes and void fraction, which is the volume of porous space in relation to the overall volume (volume of porous space plus that of microparticles). In reports on granular hydrogels, a wide range of these properties has been reported but generally, porosity is cellular scale (10-50 µm), and void fractions are in the range of 0.1 to 0.6. Here, the average void fraction of our PEDOT:PSS granular hydrogel was 0.19. The average pore diameter was 25.84 µm and, while the observed range was 9.05 µm to 67.47 µm in diameter, 96.66% of pore diameters fell within the range that has been shown to support cells among them (10-50 µm, Supplementary Figure 5).

**Figure 5.**
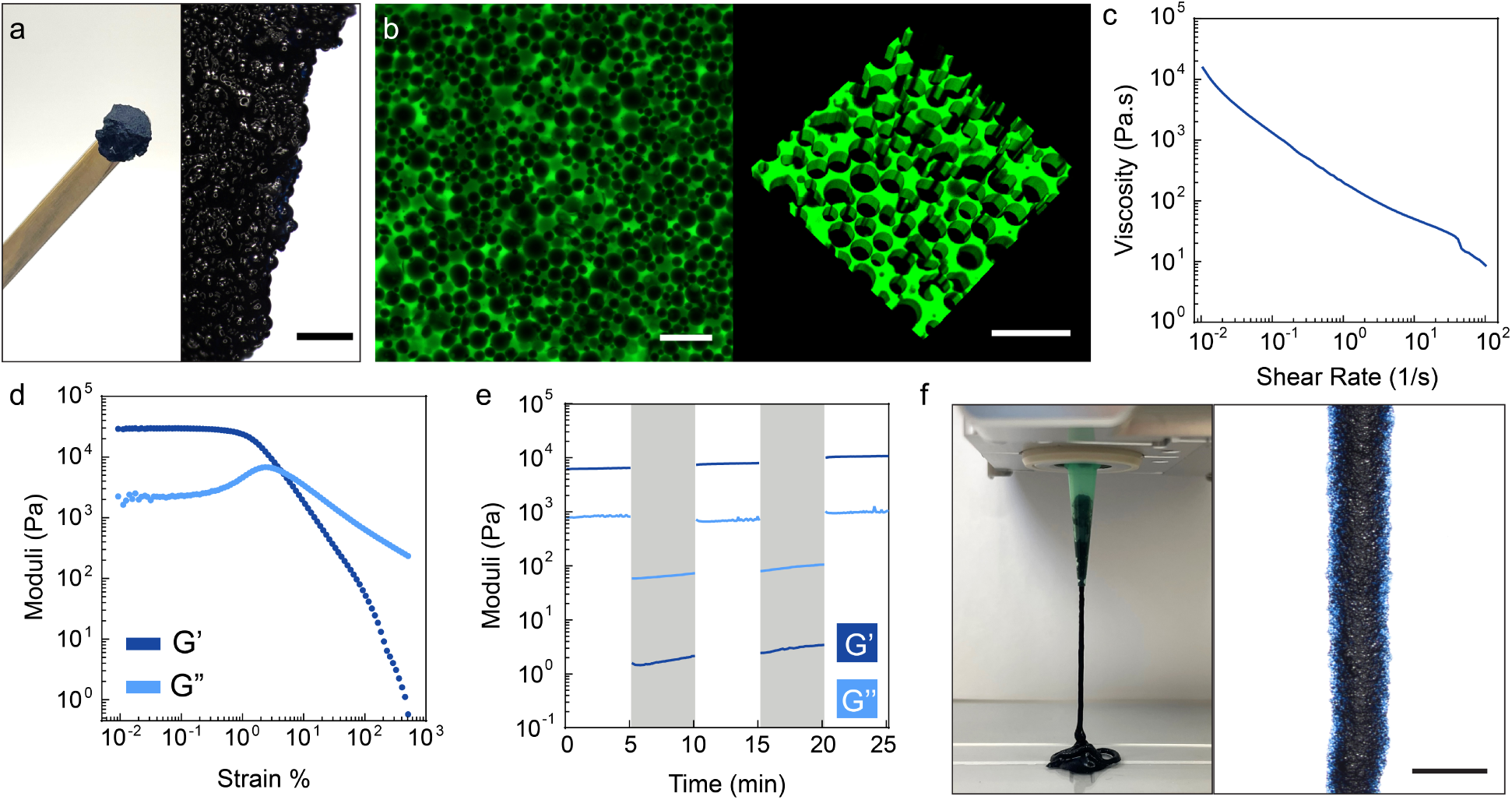
Densely packed PEDOT:PSS microparticles form a granular hydrogel as evidenced by interconnected microporosity and dynamic mechanical properties that facilitate extrusion via 3D printing. A) Microparticles were densely packed with vacuum filtration, resulting in a paste-like material that could be manipulated with a microspatula. Scale bar for right image 500 µm. B) Image from an inverted epifluorescence microscope (left) and 3D rendering of Z stack (68 µm in depth) from confocal microscopy (right). Interconnected microporosity is evident from fluorescent voids between densely packed microparticles. Scale bar 100 µm left, 200 µm right. C-E) Oscillatory shear and rotational rheology of PEDOT:PSS granular hydrogels. C) Shear-thinning behavior of the densely packed microparticles was observed as evident by a decrease in viscosity in response to increasing shear rate. D) The material exhibited deformation and underwent a gel-to-liquid transition at ∼5% strain (1 Hz). E) A time sweep with alternating applications of low (unshaded, 0.5%) and high strain (shaded, 500%) indicated recovery of the material (G’>G”) at low strains following high strain (1 Hz). F) Extrusion of the granular hydrogel through an 18 G nozzle achieved with pneumatic 3D printing. Scale bar of right image 5 mm.

To further confirm the successful generation of a granular material, the dynamic mechanical properties of the densely packed PEDOT:PSS microparticles were investigated by oscillatory shear rheology. The densely packed microparticles were loaded onto the Peltier plate and, under the 20 mm parallel plate geometry, formed a disc that others have identified as optimal for granular hydrogel rheology^54^(Supplementary Figure 6). During an angular frequency sweep, the elastic response was independent of frequency between 0.1 and 10 Hz (Supplementary Figure 7). Therefore, a frequency of 1 Hz was used in all further testing. When the microparticles were subjected to an increasing shear rate, the viscosity decreased, confirming the material is shear-thinning (Figure 5c). In a strain sweep, at small strains within the linear region, the storage modulus was higher than the loss modulus indicating dominant solid-like, elastic behavior (Figure 5d). As the percent strain increased, the material entered the nonlinear region, exhibited signs of deformation, and underwent a gel-to-liquid transition as shown by the storage modulus and loss modulus crossover. Next, the self-healing capabilities of the material were investigated under conditions of 0.5% low strain and 500% high strain (as identified by the strain sweep). In the first cycle of 0.5% and 500% strain, the material exhibited expected behavior as indicated by the strain sweep (gel at low strains, liquid at high strains) (Figure 5e). The deformation observed at the high strain was shown to be recoverable as evidenced by restoration of previous storage moduli values within 0.1 seconds of when the percent strain was lowered. Collectively, these behaviors are attributed to shear stress overcoming load-bearing chains. Contacts between microparticles are broken and distances between microparticles widen, leading to a decrease in viscosity and shear-yielding to liquid-like behavior. When shear stress is lessened or removed, the movement of the microparticles is greatly reduced, allowing the microparticles to make stable contact again, significant interparticle interactions to resume, and gel characteristics to return. In summary, the presence of both dynamic mechanical properties and the interconnected microporous structure confirm that the densely packed PEDOT:PSS microparticles form a granular hydrogel.

Finally, to determine if these dynamic mechanical properties give rise to application-desirable features such as extrudability for 3D printing or injection, we tested extrusion of the granular hydrogel. Shear-yielding and shear-thinning are expected to facilitate extrusion while self-healing subsequently facilitates the maintenance of structure’s shape^13^. We performed extrusion by 3D printing the granular hydrogel using 16 (∼1194 µm inner diameter), 18 (∼838 µm inner diameter), and 20 gauge (∼584 µm inner diameter) tapered tips (Figure 5f, Supplementary Figure 8a). Printed lines were achieved and the average width of the resulting 3D printed strands decreased with decreasing nozzle diameters (Supplementary Figure 8b). The ability to extrude and maintain 3D printed structures shows promise for future applications dependent on extrudability and maintenance of extruded structures as already demonstrated in non-conducting granular hydrogels ^20,55–57^.

### PSS Removal from Microparticles Enhances Granular Hydrogel Conductivity

Due to our selection of an intrinsically conducting polymer, electronic conductivity is expected *within* individual microparticles among the hydrogel network. However, on the millimeter scale, conductivity of the granular material requires conductivity *between* microparticles. Electronic conductivity among conducting particles is achieved when particles are in contact, such as in composites where high particle densities are used to achieve percolating networks^58^. Given that we have already achieved dense particle packing as it is an inherent characteristic that defines granular hydrogels and their mechanical behaviors, we expected this particle packing to be sufficient for achieving conductivity, as others have shown^20,23^.

When densely packed by vacuum filtration, we found that the PEDOT:PSS granular hydrogel achieved an average conductivity of 7.128 ± 6.536 S/m. This conductivity exceeded that reported of most composite granular hydrogel materials (∼10^-4^ to 10^0^ S/m)^20,21^ and was similar to the conductivity reported for a PEDOT:PSS granular hydrogel made through mechanical fragmentation (10.8 S/m)^22^. However, the conductivity values were inconsistent (91.70% coefficient of variation) which could prove problematic for the material’s use in bioelectronic devices. Therefore, we sought to address this by producing microparticle populations with more consistent conductivity.

Aqueous dispersions of PEDOT:PSS consist of nano-scale aggregates of this polymer complex. PSS and PEDOT are bound together by electrostatic interactions between the negatively charged PSS and positively charged PEDOT chains. Commercially available PEDOT:PSS colloidal dispersions are often made with an excess of PSS to PEDOT (approximately 2.5:1 weight ratio or 1.9 molar ratio of PSS to PEDOT for Heraeus Clevios™ PH 1000 used here)^59^. The presence of PSS is two-fold: one purpose is to serve as a dopant, while the other is to impart aqueous stability and solution processability to the colloidal dispersion.

However, PSS itself is insulating. When formed from these colloidal dispersions, materials are characterized by PEDOT-rich domains surrounded by PSS-rich domains. To increase conductivity, materials are processed to remove excess PSS to grow PEDOT-rich regions and bring these regions closer together^60,61^. The conductivity of PEDOT:PSS materials is commonly increased by (1) incorporation of ion pairs or co-solvents in the casting dispersion that are later often removed from the material by washing or (2) post-fabrication treatment of materials through immersion of the material in acids or polar solvents, hereafter referred to as “post-treatment”. These methods work to facilitate PSS removal by inducing charge screening of PEDOT and PSS^60,62,63^.

We have previously demonstrated that increasing amounts of ionic liquid (IL) added to PEDOT:PSS colloidal dispersions result in hydrogels of increasing conductivity^34^. Additionally, we have applied post-treatment to fabricated hydrogels to increase conductivity^35^. Towards our objective of improving conductivity of the generated granular hydrogels, we selected one of each of these methods: incorporation of ionic liquid into the PEDOT:PSS colloidal aqueous dispersion to fabricate microparticles and additional post-treatment of fabricated microparticles with 17.3 M acetic acid (Supplementary Figure 9). It should be noted that these treatments are not expected to participate in the hydrogel polymer network, as we perform significant washing that prior studies have shown to result in their removal^64,65^. Therefore, the nomenclature of groups investigated here (PEDOT:PSS/IL, PEDOT:PSS/AA, PEDOT:PSS/IL/AA) refers to their fabrication methods, not their composition as all groups are expected to be solely comprised of conducting polymer.

Ionic liquid was added to the PEDOT:PSS colloidal dispersion at a concentration of 10 mg mL^-1^. Upon subsequent addition to a 90 °C oil phase, PEDOT:PSS/IL microdroplets were formed. Removal from the oil phase and suspension in phosphate-buffered saline (PBS) confirmed fabrication of aqueous stable microparticles (Figure 6a). To investigate post-treatment, PEDOT:PSS and PEDOT:PSS/IL microparticles were immersed in 17.3 M acetic acid overnight. When resuspended in PBS, acetic acid-treated PEDOT:PSS (PEDOT:PSS/AA) and acetic acid-treated PEDOT:PSS/IL microparticles (PEDOT:PSS/IL/AA) were both aqueous stable. PEDOT:PSS/IL/AA microparticles appeared consistent in color (Figure 6a); however, PEDOT:PSS/AA microparticles appeared heterogeneous in color darkness, while some appeared translucent (Supplementary Figure 10). These changes could be indicative of hydrogel material loss and thus, this group was not investigated further. In addition to optical microscopy, scanning electron microscopy (SEM) also indicated that there were little to no obvious differences between the microparticle populations (Figure 6c). All particle groups exhibited a spherical shape and evidence of a rough surface, although this cannot be concluded given the processing of lyophilization needed for visualization by this method.

**Figure 6.**
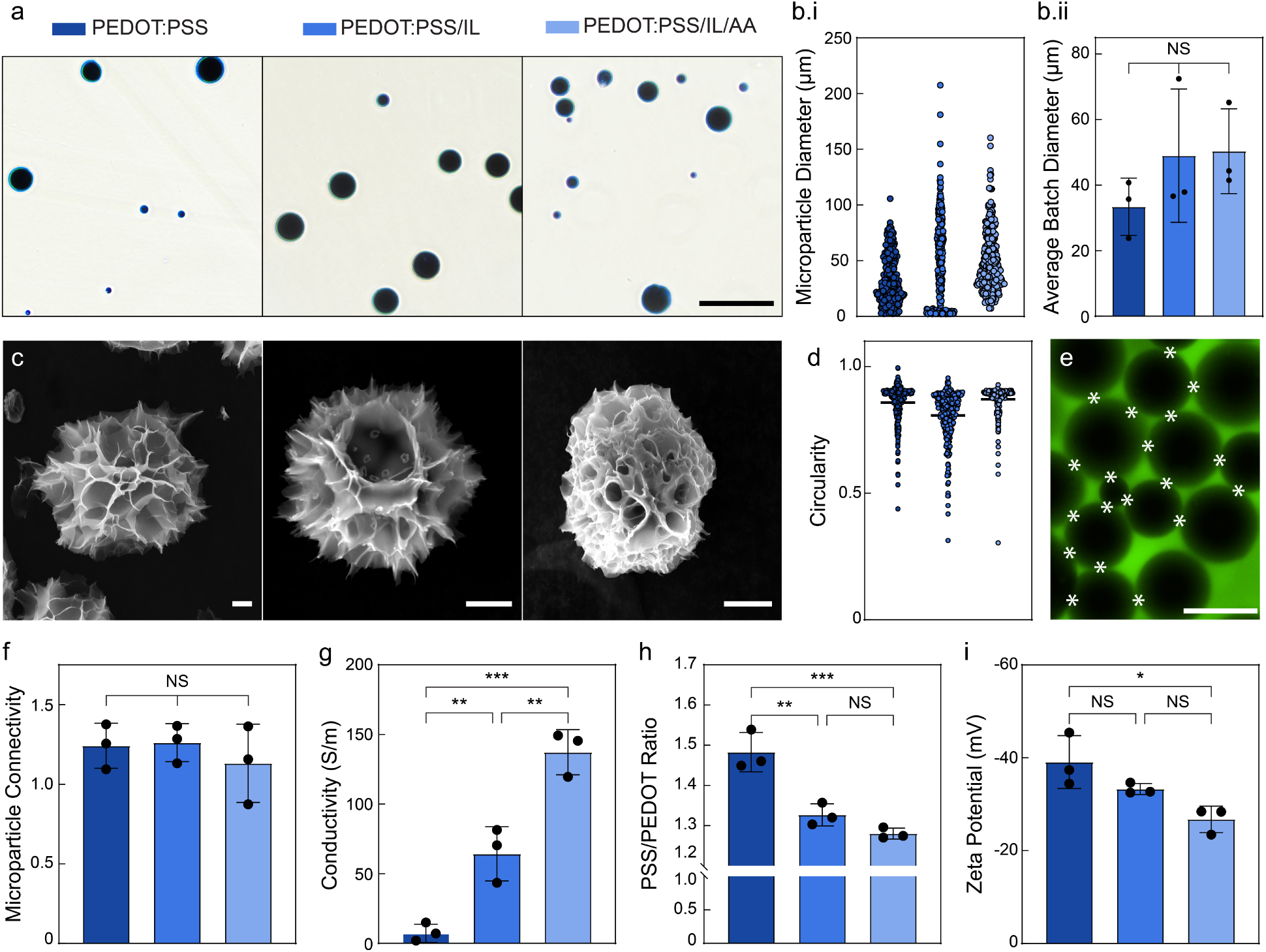
Ionic liquid incorporation into PEDOT:PSS colloidal dispersion and post-fabrication treatment with acetic acid facilitate PSS removal from microparticles for enhanced granular hydrogel conductivity. PEDOT:PSS microparticles formed by adding IL to the colloidal aqueous dispersion prior to injection into oil bath. Further post-treatment of fabricated microparticles with 17.3 M acetic acid was also performed. Population analysis of PEDOT:PSS microparticles at 90 °C is presented here again from Figure 4 (representative image and diameter and circularity values) to facilitate comparison with these microparticle groups (PEDOT:PSS/IL and PEDOT:PSS/IL/AA). A) PEDOT:PSS/IL and PEDOT:PSS/IL/AA microparticles were aqueous stable and similar in appearance to PEDOT:PSS microparticles as observed by optical microscopy (particles in PBS). Scale bar 200 µm. Bi) Diameters of microparticle population (N=300 from 3 fabrication batches, 100 microparticles quantified per batch). Bii) Mean diameter per fabrication batch (N=3). Fabrication batch values obtained by averaging values of 100 analyzed microparticles. Standard deviation presented. C) Scanning electron micrographs showed all conditions resulted in spherical microparticles with similar topography. Scale bar 15 µm. D) Circularity values of microparticle population (N=300 from 3 fabrication batches, 100 microparticles quantified per batch, line denotes mean). All groups resulted in spherical microparticles with mean circularity values exceeding 0.80. E) Suspending each microparticle population in FITC-Dextran aqueous solution followed by vacuum filtration facilitated visualization of the pores among microparticles of granular hydrogels. The connectivity of densely packed microparticles was evaluated by counting the number of microparticle-microparticle junctions (defined as discrete points of contact between two adjacent microparticles, white asterisks) and normalizing to the number of microparticles in the sample. Scale bar 50 µm. F) When densely packed to form granular hydrogels, the number of microparticle-microparticle junctions normalized to the number of microparticles in the sample showed no significant differences between groups. G) Conductivity of the granular hydrogel increases when comprised of PEDOT:PSS/IL and PEDOT:PSS/IL/AA microparticles in comparison to PEDOT:PSS microparticles (7.128 S/m, 64.25 S/m, and 137.2 S/m for PEDOT:PSS, PEDOT:PSS/IL, and PEDOT:PSS/IL/AA, respectively). H) The PSS/PEDOT ratio quantified with XPS showed a reduction from 1.483 in PEDOT:PSS microparticles to 1.327 in PEDOT:PSS/IL microparticles to 1.280 in PEDOT:PSS/IL/AA microparticles. I) Zeta potential measurements indicated less negative surface potential with IL incorporation and AA post-treatment. Mean and standard deviation presented for N=3 samples (F,G, H, I). One-way analysis of variance (ANOVA) and Tukey’s multiple comparison test. ****P ≤ 0.0001 ***P ≤ 0.001 **P ≤ 0.01 *P ≤ 0.05 non-significant (NS) P > 0.05.

To quantify potential effects of these treatments on the microparticle size and shape, both of which are important for granular hydrogel formation, we analyzed the diameters and circularity of the various populations. In comparison to the average batch diameter of PEDOT:PSS microparticles (33 µm), the average batch diameter of PEDOT:PSS/IL and PEDOT:PSS/IL/AA microparticles increased to 49 µm and 50 µm, respectively (Non-significant (NS), Figure 6b.ii). In analysis of the microparticle population, the minimum individual microparticle diameters recorded were similar for PEDOT:PSS and PEDOT:PSS/IL (2 µm) but PEDOT:PSS/IL/AA (7 µm) was slightly larger (Figure 6b.i). This group required a new filtration mesh with 10 µm pores as part of the post-treatment process (for particle isolation from acetic acid, see Methods) that likely caused loss of smaller particles. The maximum individual microparticle diameters were higher for PEDOT:PSS/IL and PEDOT:PSS/IL/AA, but these larger particles (>100 µm) comprise very little of the entire population (less than 10%). If more defined diameter characteristics are needed, filtration with meshes of varying pore sizes can be used to refine the population. For example, a sample of PEDOT:PSS/IL/AA microparticles passed through a cell strainer (PluriSelect) with 60 µm diameter pores reduced the maximum diameter from 120 µm to 54 µm (Supplementary Figure 11). In all, the size ranges without further selection are still compatible with guidelines for generation of granular hydrogels, which is the dense packing of microparticles with diameters of tens to hundreds of micrometers^55,66^. Additionally, when compared to the starting population of PEDOT:PSS microparticles, PEDOT:PSS/IL and PEDOT:PSS/IL/AA had similar ranges of circularity values (∼0.30 to ∼0.96, Figure 6d). Lower circularity values (circularity value < 0.70) correspond to PEDOT:PSS fragments generated during the batch emulsion fabrication process but only constitute 15% or less of each microparticle population. The majority of the microparticles exhibit a fairly spherical shape as showcased by the average circularity being greater than 0.8 (0.8575, 0.8103, and 0.8711 for PEDOT:PSS, PEDOT:PSS/IL, and PEDOT:PSS/IL/AA respectively; Figure 6d). In summary, either of these methods (IL incorporation, acetic acid treatment) result in size characteristics that should be compatible with granular hydrogel formation and shape characteristics favored for extrusion and microporosity.

Next, we confirmed granular hydrogel formation by packing particles of these two methods and performing oscillatory shear rheology. Both the PEDOT:PSS/IL and PEDOT:PSS/IL/AA granular hydrogels possessed expected shear-thinning and self-healing dynamic mechanical properties, similar to that of the PEDOT:PSS granular hydrogel (Supplementary Figure 12). Before evaluating our initial objective of increasing conductivity of the granular hydrogel, we first wanted to confirm that these granular hydrogels comprised of particles of various treatments resulted in similar particle packing. Particle packing is important for us to characterize for interpreting our conductivity results since granular hydrogel conductivity is expected to be dependent on both particle packing (networks of contacting microparticles) as well as the conductivity of individual particles themselves. We evaluated microparticle connectivity by quantifying the number of microparticle-microparticle junctions (defined as discrete points of contact between two adjacent microparticles) and normalizing that quantity to the number of microparticles present in the granular hydrogels (Figure 6e). There were no significant differences in the normalized microparticle-microparticle junctions indicating there were no differences in particle packing between the different granular hydrogels (Figure 6f). Therefore, any observed differences in conductivity should be due solely to the changes in the particles themselves.

Next, a four-point probe was used to measure the conductivity of the granular hydrogels. In comparison to that of the PEDOT:PSS granular hydrogel (7.128 S/m), the PEDOT:PSS/IL and PEDOT:PSS/IL/AA granular hydrogels possessed a significantly greater conductivity of 64.25 and 137.2 S/m, respectively (801.4% and 1824% increase to the PEDOT:PSS granular hydrogel, respectively; Figure 6g). Additionally, conductivity measurements of these groups become more consistent (coefficients of variation: PEDOT:PSS 91.70%, PEDOT:PSS/IL 30.38%, PEDOT:PSS/IL/AA 11.81%). These conductivity values of the granular hydrogels fall within similar ranges reported (1-100 S/m) for bulk conducting hydrogels that have been successfully used as part of bioelectronic devices, such as organic electrochemical transistors and skin electrodes for electrophysiological monitoring^67–69^.

To confirm the increase in conductivity observed was due to improving conductivity of microparticles through PSS removal, we examined measures of relative contents of PSS to PEDOT. To calculate the PSS/PEDOT ratio, X-ray photoelectron spectroscopy (XPS) was performed on microparticles to examine the sulfur 2p orbital spectra. Binding energy peaks within 157-161 eV corresponded to thiophenes of PEDOT and peaks within 162-167 eV corresponded to sulfonates of PSS (Supplementary Figure 13). PEDOT:PSS microparticles generated from the colloidal dispersion alone possessed a PSS/PEDOT molar ratio of 1.483 (Figure 6h). This ratio was less than the estimated molar ratio of pristine PEDOT:PSS colloidal dispersion (∼1.9), indicating that PSS removal takes place during the microparticle fabrication process which includes water washing. When compared to PEDOT:PSS microparticles, PEDOT:PSS/IL and PEDOT:PSS/IL/AA microparticles had significantly smaller PSS/PEDOT molar ratios of 1.327 and 1.280, respectively, indicating further PSS loss. The anticipated amount of PSS removal and subsequent change in PSS/PEDOT ratio is dependent on the starting material properties and the treatment selected. For example, treatment with sulfuric acid (H_2_SO_4_) has resulted in some of the lowest PSS/PEDOT ratios reported (0.5)^59^. Post-treatment of PEDOT:PSS hydrogels with more mild treatments (dimethyl sulfoxide, ethanol, and 60% acetic acid solution) has resulted in more moderate PSS/PEDOT ratios of 1-1.35 and, thus, are similar to ratios reported here^35^.

To further examine loss of PSS, we analyzed the zeta potential of the particles. PEDOT:PSS aggregates of the starting colloidal dispersion has been reported to have a zeta potential of approximately −50 mV^60,62,70^. Given the XPS results indicating loss of negatively charged PSS that was still in excess to positively charged PEDOT, we expected a higher but still negative zeta potential. In order of their PSS/PEDOT ratios, PEDOT:PSS, PEDOT:PSS/IL, and PEDOT:PSS/IL/AA had zeta potentials of −39.08 mV, −33.29 mV, and −26.67 mV, respectively, showing a similar decreasing trend (Figure 6i). In conclusion, similar particle connectivity within the granular hydrogels as well as decreasing trends of PSS/PEDOT ratios and zeta potentials of particles themselves all support that PSS removal from PEDOT:PSS microparticles is responsible for improving granular hydrogel conductivity.

### PEDOT:PSS Granular Hydrogel as a Bioencapsulating Electrode

Cytocompatibility plays an important role in material applications such as injectables, 3D bioprinting of cell-laden materials, and 3D tissue engineering scaffolds where the microparticles will be interfacing with cells and tissues. Therefore, we aimed to establish cytocompatibility of the microparticles utilizing methods adopted from ISO 10993-5, standard test methods previously applied to evaluate hydrogel cytocompatibility^71,72^. To begin cytocompatibility assessment, normal human dermal fibroblasts were seeded in 24 well plates and allowed to grow for four days prior to the application of PEDOT:PSS, PEDOT:PSS/IL, and PEDOT:PSS/IL/AA microparticles. Cells remaining in culture without the application of microparticles served as a negative control (absence of cytotoxicity). Twenty-four hours after application, the microparticles were removed and cell viability was quantified using a LIVE/DEAD^TM^ staining kit and Hoechst (Figure 7a). According to ISO 10993-5, biomaterials are considered cytocompatible if they result in a cell viability reduction of less than 30% in comparison to the control. Here, cell viability in the control and all microparticle conditions was 98% or greater, confirming the microparticles are cytocompatible according to the ISO standard (Figure 7b). Additionally, fibroblasts were found growing around microparticles and conforming to the curvature of the material in all three conditions as indicated by the white arrows in Figure 7a and Supplementary Figure 14. Beyond cytocompatibility, we also observed indications of long-term particle stability. The different particle groups appeared aqueous stable for a minimum of 30 days as the particles did not show any visible changes indicative of material degradation (Supplementary Figure 15). The particles can also be lyophilized and re-hydrated, indicating a means to potentially store particles indefinitely prior to re-hydration before immediate use (Supplementary Figure 16). Collectively. these results are positive first steps in evaluating the conducting granular hydrogels for cell and tissue interfacing.

**Figure 7.**
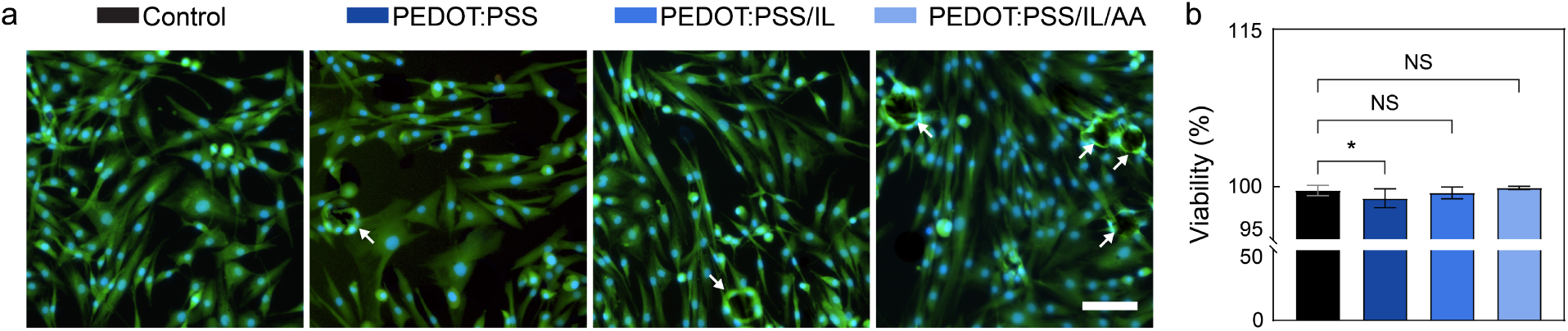
PEDOT:PSS microparticles are cytocompatible when in contact with normal human dermal fibroblasts. A) Representative epi-fluorescence images of normal human dermal fibroblasts (NHDFs). Microparticles were seeded on monolayers of NHDFs for twenty-four hours, and fibroblasts with no microparticles acted as a control. After removal of microparticles, fibroblasts were stained with LIVE/DEAD™ (live cells – Calcein AM/green; dead cells – ethidium homodimer-1/red) and Hoechst (blue nuclear stain for all cells). Fibroblasts were found to have grown around individual particles (arrows). Scale bar 100 µm. B) Mean cell viability was 98% or greater in all conditions. Mean and standard deviation presented. N=4 cell culture wells per condition. One-way analysis of variance (ANOVA) and Dunnett’s multiple comparison test. ****P ≤ 0.0001 ***P ≤ 0.001 **P ≤ 0.01 *P ≤ 0.05 non-significant (NS) P > 0.05.

Next, we investigated if the conducting granular hydrogels could be used as electrodes in electrophysiological monitoring and if the dynamic mechanical properties could be leveraged for achieving bioencapsulation, where the material envelops cells or tissue structures. In mammals, electrophysiological monitoring is used widely for studying how ionic currents are created and transmitted in cells and tissues for better understanding of various physiological processes and diseases. Researchers have also been using electrophysiological monitoring to study invertebrates^73,74^. More specifically, investigation of invertebrate olfactory systems is needed to understand the processes by which their neural circuits process olfactory information and use this information to navigate their environment such as during flight. Beyond basic biological investigations, biohybrid devices combining insect olfaction with man-made technology can create advanced sensors for complex and dynamic environments where engineered chemical sensors fall short^75^. Such biohybrid sensors could be used for detecting various chemicals towards real-time assessment for food contaminants to reduce outbreaks and food waste, more sensitive environmental monitoring of the presence of hazardous chemicals, and detection of low concentration volatile organic compounds produced from cancer cells in patients’ blood for minimally invasive and cost-effective clinical diagnostics^76,77^.

In insects, odors are detected by olfactory receptor proteins expressed by olfactory receptor neurons in the insect’s antenna. When an odor molecule binds to receptor proteins, they depolarize the cell and action potentials are triggered^78^. The summed response of all the olfactory receptor neurons present in the antenna can be measured as a change in the potential between the two ends of the antenna. This recording method is usually referred to as electroantennography (EAG)^79^. The antenna is cylindrical and therefore makes limited contact with two-dimensional planar electrodes for measurements. Instead, measurements have been procured using glass capillary electrodes filled with an electrolyte solution which contacts the antenna, chlorodized silver wires inserted directly into the antenna, as well as through encapsulation of the antenna in an electrolyte gel^79–81^. We proposed to use the conducting granular hydrogel to form encapsulating electrodes around the ends of an excised locust antenna and record potentials generated in response to odor exposures (Figure 8a).

**Figure 8.**
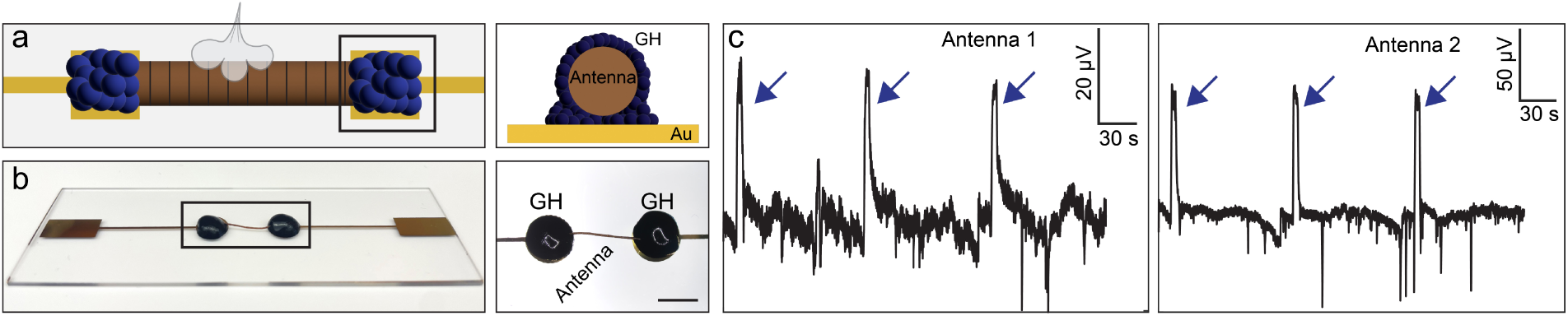
Bioencapsulating electrodes made from PEDOT:PSS granular hydrogel record potential changes in locust electroantennograms. A) Schematic of proposed set-up in which locust antenna is encapsulated within the granular hydrogel (labeled GH) placed atop gold to assemble the bioencapsulating electrode. Bioencapsulating electrodes proposed for electroantennograms, monitoring potential changes across the antenna in response to odor deliveries. B) The PEDOT:PSS/IL/AA granular hydrogel was deposited by a microspatula on top of gold pads and both ends of the antenna were encapsulated within the granular hydrogel. Scale bar of right image 5 mm. C) Potential changes across the antenna in response to delivery of hexanol detected and indicated with blue arrows.

The PEDOT:PSS/IL/AA granular hydrogel was deposited on the electrodes and the two ends of the antenna were successfully encapsulated within the two granular hydrogel deposits, so an exposed portion of the antenna spanned the distance between the two electrodes (Figure 8b). Hexanol, a green leaf volatile, was delivered three times to two antennae via a pneumatic pico-pump controlled by a custom MATLAB script. Absolute values of amplitudes around 30 and 90 µV were recorded upon odor application in all three trials for two antennae (Figure 8c). While the absolute value of the extracellular potential can vary across preparation of the antenna, the signal change arising due to odor exposure was clearly detectable with high signal-to-noise ratio. These successful recordings that are comparable with those found in literature validate our claim that the conducting granular hydrogel can act as a bioencapsulating electrode in electrophysiological monitoring applications^80^.

## Conclusion

In summary, we have reported novel methodology to fabricate spherical, pure PEDOT:PSS microparticles for generating conducting granular hydrogels. First, 90 °C was established as the optimal of the studied temperatures for producing smaller and more uniformly shaped, aqueous stable PEDOT:PSS microparticles. Vacuum filtration achieved dense packing of the microparticles and resulted in successful granular hydrogel formation, as evident by a paste-like material with an interconnected microporous structure. Rheological investigations of the PEDOT:PSS granular hydrogel confirmed shear-thinning and self-healing properties. Extrudability of the PEDOT:PSS granular hydrogel and maintenance of 3D printed strands were also achieved. PEDOT:PSS granular hydrogels were found to be conducting but measurements were inconsistent. We increased and obtained more consistent granular hydrogel conductivity by fabricating particles with lower PSS/PEDOT ratios, as confirmed by XPS and zeta potential. These particles were achieved by applying methods previously reported on PEDOT:PSS materials to facilitate PSS removal and included inclusion of ionic liquid in the PEDOT:PSS colloidal dispersion added to the oil phase and further post-treatment of the resulting microparticles with acetic acid. These methods resulted in a maximum conductivity value of 137 S/m and a reduced coefficient of variation (11.81%). These methods had no impact on granular hydrogel formation as confirmed by maintenance of dynamic mechanical properties via oscillatory rheology. Exposure of human dermal fibroblasts to the various microparticles for 24 hours showed cytocompatibility with all conditions resulting in high viability (>98%) and fibroblasts growing around microparticles.

We envision these conducting granular hydrogels could be utilized as 3D printed customized electrodes that can conform to topographically diverse surfaces or completely encapsulate biological components^33,65,82^, 3D tissue engineering scaffolds that can monitor and/or stimulate encapsulated cell viability^83^, as well as injectable therapies that can facilitate transfer of electrophysiological signals through gaps in the tissue while facilitating cell recruitment for tissue repair^84^. To demonstrate the ability of the conducting granular hydrogel to act within these capacities, we applied the PEDOT:PSS granular hydrogel as a bioencapsulating electrode for electroantennograms of insect antennae. The conducting granular hydrogel was used to successfully monitor potential changes generated across locust antennae upon exposure to an odor. Further study and optimization will be required to realize the full capabilities of the material, but our results demonstrate that the material holds great promise for achieving new capabilities in bioelectronics.

## Methods

### Fabrication of PEDOT:PSS Microparticles with Water-in-Oil Emulsion

Approximately 700 mL of food grade mineral oil with 0.05% (wt%) sorbitane monooleate (Span®-80, Sigma Aldrich) was stirred at 100 RPM and allowed to equilibrate at the relevant temperature for 60 minutes. Unless otherwise stated, the oil phase was held at 90 °C. Colloidal aqueous dispersions of PEDOT:PSS (Clevios PH 1000) were acquired from Heraeus. Certificates of analysis provided by the supplier show the colloidal aqueous dispersions used in this study ranged from 1.0 to 1.3 wt%. PEDOT:PSS dispersion was vortexed for one minute prior to use. Ionic liquid (IL), 4-(3-butyl-1-imidazolio)-1-butanesulfonic acid triflate (Santa Cruz Biotechnology), was prepared at a concentration of 100 mg mL^-1^ in 0.01 M hydrochloric acid prior to addition to PEDOT:PSS.

For fabrication of microparticles, 1 mL of the PEDOT:PSS dispersion alone or PEDOT:PSS dispersion with 10 mg mL^-1^ IL was injected via a syringe with a 32G blunt needle into an oil bath stirred at 750 RPM (Supplementary Video 1). Constant pressure was applied to the syringe plunger to achieve one continuous stream. The temperature and stir speed were maintained for one hour prior to removal from heat and subsequent washing to remove the oil phase.

For post-treatment of microparticles, microparticles were immersed in acetic acid (17.3 M) at a concentration of 5-10 mg of microparticles mL^-1^.

### PEDOT:PSS Microparticle Washing

Once cooled to room temperature, the microparticles were removed from the mineral oil using either vacuum filtration or oil phase aspiration. Vacuum filtration was conducted by pouring the microparticles into a Buchner funnel with 90 mm diameter cellulose filter paper (Whatman) under a vacuum. Isolated particles were collected by rinsing the filter paper with deionized water into a collecting container. To facilitate oil aspiration, deionized water was poured into the emulsion and allowed to settle for 20 minutes. Due to differences in density, the microparticles and water settled at the bottom of the beaker forming an aqueous layer, while the mineral oil, being less dense, rose to form a separate layer on top. The top layer of mineral oil was aspirated without disturbing the aqueous layer before pipetting the aqueous layer into a collecting container, conical centrifuge tubes. To remove mineral oil remnants, phosphate buffered saline (PBS: Sigma Aldrich P3813) was added to microparticles to reach near the top of the tube and sonicated for 10-15 minutes. To perform repeated washing, microparticles were packed by centrifugation at 5400 RCF for 10 minutes to facilitate aspiration and PBS exchange. A small amount (1-5 mL) of deionized water was used to resuspend microparticles in the remaining PBS after aspiration and transfer the suspension to a new centrifuge tube at every PBS exchange. PBS was then added to the new centrifuge tube and the washing steps of sonication, centrifugation, and PBS replacement were completed three additional times. Removal of the oil phase was confirmed by visually inspecting the particle suspension with a stereoscope (Nikon SMZ1270) for the absence of mineral oil droplets.

For acetic acid treated particles, the microparticles were vacuum filtered from the acetic acid using a cell strainer with a 10 µm mesh (pluriStrainer, PluriSelect). Microparticles were then resuspended in and washed with four one-hour exchanges of deionized water followed by one overnight wash. To perform washes, microparticles were packed by centrifugation at 5400 RCF for 10 minutes to facilitate aspiration and water exchange. Microparticle suspensions were agitated on an orbital shaker at 200 RPM during washes.

### PEDOT:PSS Aqueous Phase and Microparticle Size Measurements

Aqueous phase droplets in the oil phase were imaged using a stereoscope (Nikon SMZ1270) and diameters were measured with NIS-Elements (Nikon). Once isolated from the oil phase and suspended in water, PEDOT:PSS microparticles were imaged using a Nikon Eclipse Ts2-FL inverted microscope. Particle diameters were measured using image analysis with a custom MATLAB script. Microparticle circularity was measured using image analysis with a custom ImageJ script. Analysis of microparticle diameter and circularity values was conducted on 3 fabrication batches of microparticles, with 100 microparticles sampled from each batch to analyze the population. Mean values of diameter and circularity were presented by averaging the measurements of the 100 sampled microparticles in each batch.

### PEDOT:PSS Colloidal Dispersion and Granular Hydrogel Rheology

The viscoelastic behavior and gelation kinetics of the PEDOT:PSS colloidal aqueous dispersion were characterized with a TA Instruments HR-20 rheometer. Measurements were performed using a cone-plate fixture (stainless steel plate at 40 mm diameter and 1.98889° cone geometry) with a solvent trap. Unless otherwise stated, oscillatory tests on the PEDOT:PSS colloidal aqueous dispersion were performed at 1% strain and 1 rad s^-1^ angular frequency.

The rheological properties of the granular hydrogels were quantitatively evaluated using a TA Instruments HR-20 rheometer. All measurements were taken with a 20 mm parallel plate geometry set at a gap height of 1.1 mm at 25 °C. The densely packed microparticles were loaded onto the Peltier plate, and, when the geometry was lowered to a gap height of 1.1 mm, formed a disc (Supplementary Figure 6). A frequency sweep at 1% strain was conducted to quantify at which frequencies the material would exhibit a linear trend and identify a frequency at which to conduct subsequent tests (Supplementary Figure 7). Unless otherwise stated, all oscillatory tests on granular hydrogels were performed at a frequency of 1 Hz.

### Scanning Electron Microscopy (SEM)

A Zeiss EVO 10 scanning electron microscope was used to image microparticles. Microparticles were densely packed from suspensions with vacuum filtration and transferred with a spatula to tubes, frozen at −20 °C, and lyophilized. The lyophilized samples were then mounted on stubs using carbon tape for imaging. All SEM images were captured at 20 kV with a working distance ranging from 8.5 to 9.5 mm. Brightness and contrast settings of images were adjusted to improve visibility of features.

### X-Ray Photoelectron Spectroscopy (XPS)

Microparticles were frozen at −20 °C, lyophilized, and a microspatula tip-worth of microparticles were loaded onto the XPS stub. XPS spectra were obtained with an X-ray photoelectron spectrometer (PHI 5000 VersaProbe II, Physical Electronics) according to methods described in Okafor, et al. (2025)^35^. Briefly, the C 1s and S 2p spectra were recorded with a binding energy step of 0.05 eV. Spectra peaks were calibrated using the binding energy of C 1s (284.8 eV) as a reference, and the XPS spectra of S 2p were fitted with an asymmetric line shape to investigate the properties of PEDOT:PSS. The areal ratio of the S 2p_1/2_ and S 2p_3/2_ doublet peaks were fixed at 1:2 with the binding energy splitting of each doublet peak being 1.2 eV. The S 2p doublet peak area corresponding to PEDOT and PSS was used to calculate the PSS-to-PEDOT molar ratio. Three samples were measured per microparticle condition, and one scan was taken per sample.

### Zeta Potential Measurement

Zeta potential was measured using a Malvern Panalytical Zetasizer Pro fitted with a 633 nm laser. Microparticle suspensions were analyzed at a concentration of 10 mg particles mL^-1^ in deionized water. 750 µL of each suspension was added to a folded capillary cell with gold electrodes. The analyzer, using Zetasizer Software Ultra, measured three zeta potentials via electrophoretic light scattering with mixed mode measurement-phase analysis light scattering and calculated zeta potential values using the electrophoretic mobility and Henry’s function. These three values were averaged to represent the zeta potential of each measured suspension.

### Preparation of Granular Hydrogel and Microporous Structure Analysis

Granular hydrogels were produced by achieving dense packing of PEDOT:PSS particles. This jammed state was achieved by vacuum filtrating microparticle suspensions. A cell strainer with a 10 µm mesh (pluriStrainer, PluriSelect) and a syringe in combination with a connector ring was used to create a vacuum and remove a large portion of the aqueous phase. A microspatula was then used to collect and manipulate the granular hydrogel.

To analyze microporosity of the granular hydrogel, vacuum filtered microparticles were resuspended in 10 mL of fluorescein isothiocyanate-dextran in 1X PBS (5 mg mL^-1^). Particles were then again densely packed by vacuum filtration (cell strainer set-up described above) to achieve the jammed state. A microspatula was used to fill a 4 mm diameter polydimethylsiloxane (PDMS) cylindrical mold with the granular hydrogel. The well was sealed with a glass coverslip, and the sample was then imaged using a Nikon Eclipse Ts2-FL inverted microscope. Samples were also imaged on a Leice SP8 Lightning confocal microscope and the Leica LASX 3D Analysis suite was utilized to produce 3D renderings.

The Leica LASX 3D Analysis suite was used to process images and obtain the void fraction and pore diameter measurements. The 3D Analysis Wizard was used to measure the total volume of the 3D rendering and the volume of the fluorescently labeled pores. Void fraction was calculated by dividing the volume of the fluorescently labeled pores by the total volume of the 3D rendering. The z-axes of the confocal stacks were then cropped to 15 µm to 20 µm to isolate one layer of pores. The 3D interactive measurement tool was used to measure the diameter of pores (distances between microparticles that are adjacent but non-contacting) on a singular x-y plane.

NIS-Elements (Nikon) was used to manually count microparticle-microparticle junctions (defined as discrete points of contact between adjacent microparticles) and the number of microparticles in each ROI. The normalized value of microparticle-microparticle junctions was calculated by dividing the number of microparticle-microparticle junctions by the number of microparticles.

### Conductivity

The granular hydrogel was confined to a 0.8 mm thick, 6 mm diameter polydimethylsiloxane (PDMS) cylindrical mold on top of a glass slide. The conductivity of the granular hydrogel was measured at room temperature using a four point probe (Ossila, UK) with rounded tips. The probes were positioned to be at the center of the granular hydrogel and the stage was raised until all four probes had contacted the material surface. Conductivity was determined using the Ossila Sheet Resistance Software. Conductivity values presented were means obtained from 50 recordings per measurement per sample. All measurements were procured between four and five minutes post-vacuum filtration to prevent sample dehydration.

### Cytotoxicity Study

Microparticles were isolated from 1X PBS using a cell strainer with a 10 µm mesh (pluriStrainer, PluriSelect). The microparticles were suspended in 70% ethanol at a 5-10 mg mL^-1^ overnight to disinfect the material. The microparticles were isolated from the 70% ethanol using a sterile cell strainer with a 10 µm mesh and resuspended in sterile 1X PBS at 5-10 mg mL^-1^.

Cytocompatibility was assessed according to ISO 10993-5. This ISO test dictates a material is considered cytocompatible if reduction in cell viability is less than 30% compared to that of a negative control after twenty-four hours of contact when the biomaterial contacts 10% of the established cell monolayer^85^. Normal adult human dermal fibroblasts (NHDF; Lonza CC-2511) were cultured according to supplier instructions (37 °C, 5% CO_2_). Passage 5 NHDFS were seeded at 3500 cell cm^-2^ into tissue culture treated 24 well plates and grown in FGM^TM^-2 Fibroblast Growth Medium-2 BulletKit. On day four of culture, microparticles were suspended in media and added to four wells each at 6.5 mg mL^-1^ to cover approximately 10% of the cell monolayer. Media alone served as a negative control. The cells were cultured with the microparticles for an additional 24 hours prior to cell viability analysis.

To evaluate cell viability, the microparticle-media suspensions were gently pipetted out of the wells, and the cell monolayers were fluorescently stained at room temperature for 20 minutes. The cells were stained with 2 µM Calcein AM, 2 µM Ethidium homodimer-1 (LIVE/DEAD^TM^, Invitrogen^TM^) and 10 µg mL^-1^ Hoechst 33342 (BD) in 1X PBS. Epi-fluorescence imaging was conducted using a Nikon Eclipse Ts2-FL inverted microscope. Two regions of interest (ROIs) were imaged per well at 4X magnification, and a representative image was procured of each well at 10X magnification. The images acquired at 4X magnification were analyzed using NIS-Elements (Nikon). Cell viability of each ROI was calculated by subtracting the number of dead cells from the total number of cells to determine the number of live cells. The Automatic Measurement and Thresholding functions of NIS-Elements were used to count the Hoechst-stained nuclei in the blue channel (total cells) and the ethidium homodimer-1-positive nuclei in the red channel (dead cells). The viability measurements of the microparticle groups were obtained as averages of each ROI measurement.

### 3D Printing of Granular Hydrogel

Granular hydrogel was extruded and 3D printed using a R-GEN 100 3D Bioprinter (RegenHu, Switzerland) outfitted with a pneumatic strand dispenser (pressure driven piston extrusion). Granular hydrogels formed using vacuum filtration were loaded into 3 mL syringes with gas-tight pistons for pneumatic cartridges (RegenHu) using a microspatula. The software Shaper (RegenHu, Switzerland) was used to create 3D printing strand designs. Printability assessments of the granular hydrogel through various polypropylene tapered nozzles (Nordson EFD) were conducted by extruding 15-25 mm strands at 15-25 kPa. Strands were imaged with a stereoscope (Nikon SMZ1270) and the width of each strand was measured at nine points each 2 mm apart using NIS-Elements (Nikon). These widths were averaged for each strand.

### Electroantennogram Recordings

Gold pads and connections were patterned with a laser cut mask of 5 mil Mylar adhesives. Thermal evaporation (Kurt J. Lesker Company, Nano 36™) was used to deposit a 10 nm chromium layer followed by a 150 nm gold layer. Gold patterns consisted of two 6 mm diameter circles 7 mm apart on a glass substrate. Each circle was connected to a 5 mm by 9 mm (W x L) contact pad by a 0.6 mm by 17 mm (W x L) connection. Antennas were excised from two sixth-instar locusts (*S. Americana*) from a crowded colony. A microspatula was used to place the granular hydrogel formed from PEDOT:PSS/IL/AA microparticles on each gold pad, and an antenna was placed between the pads with each end of the antenna completely encapsulated by the granular hydrogel. Hexanol (1% v/v in paraffin oil) was introduced into a desiccated air stream (0.75 L/min) via a pneumatic pico-pump (WPI Inc., PV-820). The odor was automated and triggered over three trials using a custom MATLAB code. In each trial, hexanol was delivered for 4 s with a 60 s inter-trial interval. Signals were amplified with a gain of 100 and filtered between 0 and 50 Hz (Brownlee Precision Model 440, USA). A custom MATLAB program was used to acquire signals with a band pass filter of 0.01 to 30 Hz at a sampling rate of 15 kHz. Absolute values of recorded signals are presented.

### Statistical Analysis

For statistical analysis, unpaired t-tests and ordinary one-way ANOVAs with Tukey’s or Dunnett’s multiple comparison tests were performed with GraphPad Prism. Results are presented as means with both positive and negative standard deviation. Differences were determined to be statistically significant for p<0.05.

## Data Availability

The data that supports the findings of this study are available from the corresponding author upon reasonable request.

## Acknowledgements

This research was supported by the National Science Foundation through FR #2319060 and the Center for Engineering MechanoBiology (CEMB), an NSF Science and Technology Center, under grant agreement CMMI #15-48571. The authors also acknowledge support from Washington University in St. Louis through the Women’s Health Technologies Collaboration Initiation Grant, Center for Regenerative Medicine Seed Grant, Ovarian Cancer Research Innovation Fund Award, and the McDonnell Center for Cellular and Molecular Neurobiology Small Grant. S.S.O. acknowledges support from the McDonnell International Scholars Academy. The authors also acknowledge assistance from staff and use of instruments from the Institute of Materials Science and Engineering as well as Department of Mechanical Engineering and Materials Science, Department of Energy, Environmental & Chemical Engineering, and Department of Biology at Washington University in St. Louis. The authors would also like to thank Dianne Duncan (Director of the Washington University in St. Louis Biology Department Imaging Facility) for technical assistance regarding imaging and Bob Anleitner (Laboratory Manager II of the Washington University in St. Louis Chemical and Environmental Analysis Facility) for technical assistance regarding zeta potential analysis.

## Author Contributions

A.P.G. and A.L.R. envisioned the study and wrote this manuscript. A.P.G., J.P., S.S.O., T.L., and A.L.R. were involved in data interpretation. A.P.G. developed microparticle emulsion methodologies, performed droplet and microparticle analysis, and performed all microscopy, electrical, and mechanical characterization of microparticles and granular hydrogels. A.P.G. designed and performed cell culture experiments and fluorescent imaging. A.P.G. also designed and fabricated custom electrodes and performed electroantennogram experiments and data interpretation. J.S.Y., Y.W., and C.J.V.E. performed extensive microparticle fabrication. J.S.Y. also performed microparticle diameter measurements using the MATLAB script and conducted microparticle connectivity analysis. L.C.F. developed the custom MATLAB script to analyze microparticle diameters. J.P. conducted XPS measurements. S.S.O. performed rheology on PEDOT:PSS solutions. T.L. performed cell counting and SEM imaging. S.C., A.D., S.S., and B.R. assisted with electroantennogram experiments and data interpretation. B.A.S. contributed to methodology development and data interpretation for rheological characterizations. C.P.O. performed microparticle diameter measurements. R.M.A. assisted with microparticle fabrication.

## Ethics Declaration

## References

1. Zhang, Y., Chen, H. & Song, Y. Wearable healthcare monitoring and therapeutic bioelectronics. Wearable Electronics 2, 18–22 (2025).

2. Varma, N. et al. Remote monitoring of cardiac implantable electronic devices and disease management. Europace 25, euad233 (2023).

3. Song, J., Jeong, H. E., Choi, A. & Kim, H. N. Monitoring of Electrophysiological Functions in Brain-on-a-Chip and Brain Organoids. Advanced NanoBiomed Research 2400052 (2024) doi:10.1002/anbr.202400052.

4. Yuk, H., Lu, B. & Zhao, X. Hydrogel bioelectronics. Chem. Soc. Rev. 48, 1642–1667 (2019).

5. Pitsalidis, C., et al. Organic Bioelectronics for In Vitro Systems. Chem. Rev. 122, 4700–4790 (2022).

6. Jiao, C. et al. Hydrogel-based soft bioelectronic interfaces and their applications. J. Mater. Chem. C 13, 2620–2645 (2025).

7. Zhang, Q., Liu, X., Duan, L. & Gao, G. Ultra-stretchable wearable strain sensors based on skin-inspired adhesive, tough and conductive hydrogels. Chemical Engineering Journal 365, 10–19 (2019).

8. Wei, H. et al. Conductive 3D Ti3C2Tx MXene-Matrigel hydrogels promote proliferation and neuronal differentiation of neural stem cells. Colloids and Surfaces B: Biointerfaces 233, 113652 (2024).

9. Wang, F. et al. 3D Printed Implantable Hydrogel Bioelectronics for Electrophysiological Monitoring and Electrical Modulation. Adv Funct Materials 34, 2314471 (2024).

10. Hsu, C.-C. et al. Increased connectivity of hiPSC-derived neural networks in multiphase granular hydrogel scaffolds. Bioactive Materials 9, 358–372 (2022).

11. Tanner, G. I., Schiltz, L., Narra, N., Figueiredo, M. L. & Qazi, T. H. Granular Hydrogels Improve Myogenic Invasion and Repair after Volumetric Muscle Loss. Adv Healthcare Materials (2024) doi:10.1002/adhm.202303576.

12. Xin, S., Chimene, D., Garza, J. E., Gaharwar, A. K. & Alge, D. L. Clickable PEG hydrogel microspheres as building blocks for 3D bioprinting. Biomater. Sci. 7, 1179–1187 (2019).

13. Highley, C. B., Song, K. H., Daly, A. C. & Burdick, J. A. Jammed Microgel Inks for 3D Printing Applications. Advanced Science 6, 1801076 (2019).

14. Xin, S. et al. Generalizing hydrogel microparticles into a new class of bioinks for extrusion bioprinting. Sci. Adv. 7, eabk3087 (2021).

15. Griffin, D. R., Weaver, W. M., Scumpia, P. O., Di Carlo, D. & Segura, T. Accelerated wound healing by injectable microporous gel scaffolds assembled from annealed building blocks. Nature Mater 14, 737–744 (2015).

16. Nih, L. R., Sideris, E., Carmichael, S. T. & Segura, T. Injection of Microporous Annealing Particle (MAP) Hydrogels in the Stroke Cavity Reduces Gliosis and Inflammation and Promotes NPC Migration to the Lesion. Advanced Materials 29, 1606471 (2017).

17. Lee, H.-P. et al. Dynamically Cross-Linked Granular Hydrogels for 3D Printing and Therapeutic Delivery. ACS Appl. Bio Mater. 6, 3683–3695 (2023).

18. Xin, S., Wyman, O. M. & Alge, D. L. Assembly of PEG Microgels into Porous Cell-Instructive 3D Scaffolds via Thiol-Ene Click Chemistry. Adv Healthcare Materials 7, 1800160 (2018).

19. Zhang, J. et al. “All-in-one” zwitterionic granular hydrogel bioink for stem cell spheroids production and 3D bioprinting. Chemical Engineering Journal 430, 132713 (2022).

20. Kim, S., Choi, H., Son, D. & Shin, M. Conductive and Adhesive Granular Alginate Hydrogels for On-Tissue Writable Bioelectronics. Gels 9, 167 (2023).

21. Casella, A. et al. Conductive Microgel Annealed Scaffolds Enhance Myogenic Potential of Myoblastic Cells. Adv Healthcare Materials 2302500 (2023) doi:10.1002/adhm.202302500.

22. Feig, V. R. et al. Conducting Polymer-Based Granular Hydrogels for Injectable 3D Cell Scaffolds. Adv Materials Technologies 6, 2100162 (2021).

23. Shin, M., Song, K. H., Burrell, J. C., Cullen, D. K. & Burdick, J. A. Injectable and Conductive Granular Hydrogels for 3D Printing and Electroactive Tissue Support. Advanced Science 6, 1901229 (2019).

24. Sheikhi, M. et al. Activation of muscle amine functional groups using eutectic mixture to enhance tissue adhesiveness of injectable, conductive and therapeutic granular hydrogel for diabetic ulcer regeneration. Biomaterials Advances 166, 214073 (2025).

25. Kusen, I. et al. Injectable conductive hydrogel electrodes for minimally invasive neural interfaces. J. Mater. Chem. B 12, 8929–8940 (2024).

26. Daly, A. C., Riley, L., Segura, T. & Burdick, J. A. Hydrogel microparticles for biomedical applications. Nat Rev Mater 5, 20–43 (2019).

27. Widener, A. E., Duraivel, S., Angelini, T. E. & Phelps, E. A. Injectable Microporous Annealed Particle Hydrogel Based on Guest–Host-Interlinked Polyethylene Glycol Maleimide Microgels. Advanced NanoBiomed Research 2, 2200030 (2022).

28. Shin, Y., Lee, H. S., Jeong, H. & Kim, D.-H. Recent advances in conductive hydrogels for soft biointegrated electronics: Materials, properties, and device applications. Wearable Electronics 1, 255–280 (2024).

29. Jivan, F. et al. Sequential Thiol–Ene and Tetrazine Click Reactions for the Polymerization and Functionalization of Hydrogel Microparticles. Biomacromolecules 17, 3516–3523 (2016).

30. Hirsch, M., Charlet, A. & Amstad, E. 3D Printing of Strong and Tough Double Network Granular Hydrogels. Adv Funct Materials 31, 2005929 (2021).

31. Muir, V. G., Qazi, T. H., Shan, J., Groll, J. & Burdick, J. A. Influence of Microgel Fabrication Technique on Granular Hydrogel Properties. ACS Biomater. Sci. Eng. 7, 4269–4281 (2021).

32. Huang, Y., Tang, L. & Jiang, Y. Chemical Strategies of Tailoring PEDOT:PSS for Bioelectronic Applications: Synthesis, Processing and Device Fabrication. CCS Chem 6, 1844–1867 (2024).

33. Yuk, H. et al. 3D printing of conducting polymers. Nat Commun 11, 1604 (2020).

34. Goestenkors, A. P. et al. Manipulation of cross-linking in PEDOT:PSS hydrogels for biointerfacing. J. Mater. Chem. B 11, 11357–11371 (2023).

35. Okafor, S. S. et al. 3D Printed Bioelectronic Scaffolds with Soft Tissue-Like Stiffness. Adv Materials Technologies 2401528 (2025) doi:10.1002/admt.202401528.

36. Serafin, A. et al. Electroconductive PEDOT nanoparticle integrated scaffolds for spinal cord tissue repair. Biomater Res 26, 63 (2022).

37. Tropp, J. et al. Conducting Polymer Nanoparticles with Intrinsic Aqueous Dispersibility for Conductive Hydrogels. Advanced Materials 36, 2306691 (2024).

38. Zheng, H., Jiang, Y., Xu, J. & Yang, Y. The characteristic properties of PEDOT nano-particle based on reversed micelle method. Sci. China Technol. Sci. 53, 2355–2362 (2010).

39. Rauer, S. B. et al. Porous PEDOT:PSS Particles and their Application as Tunable Cell Culture Substrate. Adv Materials Technologies 7, 2100836 (2022).

40. Jastrzebska-Perfect, P. et al. Mixed-conducting particulate composites for soft electronics. Sci. Adv. 6, eaaz6767 (2020).

41. Jain, K., Mehandzhiyski, A. Y., Zozoulenko, I. & Wågberg, L. PEDOT:PSS nano-particles in aqueous media: A comparative experimental and molecular dynamics study of particle size, morphology and z-potential. Journal of Colloid and Interface Science 584, 57–66 (2021).

42. Gutierrez-Fernandez, E., Ezquerra, T. A. & García-Gutiérrez, M.-C. Additive Effect on the Structure of PEDOT:PSS Dispersions and Its Correlation with the Structure and Morphology of Thin Films. Polymers (Basel) 14, 141 (2021).

43. Doshi, S. et al. Thermal Processing Creates Water-Stable PEDOT:PSS Films for Bioelectronics. Advanced Materials 37, 2415827 (2025).

44. Lu, B. et al. Pure PEDOT:PSS hydrogels. Nat Commun 10, 1043 (2019).

45. Friedel, B., Brenner, T. J. K., McNeill, C. R., Steiner, U. & Greenham, N. C. Influence of solution heating on the properties of PEDOT:PSS colloidal solutions and impact on the device performance of polymer solar cells. Organic Electronics 12, 1736–1745 (2011).

46. Mizrahi, B. et al. Microgels for efficient protein purification. Adv Mater 23, H258–262 (2011).

47. Widener, A. E., Bhatta, M., Angelini, T. E. & Phelps, E. A. Guest–host interlinked PEG-MAL granular hydrogels as an engineered cellular microenvironment. Biomater. Sci. 9, 2480–2493 (2021).

48. Paxton, R. A., Al-Jumaily, A. M. & Easteal, A. J. An experimental investigation on the development of hydrogels for optical applications. Polymer Testing 22, 371–374 (2003).

49. Jaeger, H. M., Nagel, S. R. & Behringer, R. P. The Physics of Granular Materials. Physics Today 49, 32–38 (1996).

50. Seymour, A. J., Shin, S. & Heilshorn, S. C. 3D Printing of Microgel Scaffolds with Tunable Void Fraction to Promote Cell Infiltration. Adv Healthcare Materials 10, 2100644 (2021).

51. Li, F. et al. Cartilage tissue formation through assembly of microgels containing mesenchymal stem cells. Acta Biomaterialia 77, 48–62 (2018).

52. Truong, N. F. et al. Microporous annealed particle hydrogel stiffness, void space size, and adhesion properties impact cell proliferation, cell spreading, and gene transfer. Acta Biomaterialia 94, 160–172 (2019).

53. Chang, C.-Y., Nguyen, H., Frahm, E., Kolaczyk, K. & Lin, C.-C. Triple click chemistry for crosslinking, stiffening, and annealing of gelatin-based microgels. RSC Appl. Polym. 2, 656–669 (2024).

54. Qazi, T. H., Muir, V. G. & Burdick, J. A. Methods to Characterize Granular Hydrogel Rheological Properties, Porosity, and Cell Invasion. ACS Biomater. Sci. Eng. 8, 1427–1442 (2022).

55. Daly, A. C. Granular Hydrogels in Biofabrication: Recent Advances and Future Perspectives. Adv Healthcare Materials 2301388 (2023) doi:10.1002/adhm.202301388.

56. D’Elia, A. M. et al. Injectable Granular Hydrogels Enable Avidity-Controlled Biotherapeutic Delivery. ACS Biomater. Sci. Eng. 10, 1577–1588 (2024).

57. Li, Y., Song, W., Kong, L., He, Y. & Li, H. Injectable and Microporous Microgel-Fiber Granular Hydrogel Loaded with Bioglass and siRNA for Promoting Diabetic Wound Healing. Small 20, 2309599 (2024).

58. Rahaman, M. et al. A New Insight in Determining the Percolation Threshold of Electrical Conductivity for Extrinsically Conducting Polymer Composites through Different Sigmoidal Models. Polymers 9, 527 (2017).

59. Kim, N. et al. Highly Conductive PEDOT:PSS Nanofibrils Induced by Solution-Processed Crystallization. Advanced Materials 26, 2268–2272 (2014).

60. Fan, X. et al. PEDOT:PSS for Flexible and Stretchable Electronics: Modifications, Strategies, and Applications. Advanced Science 6, 1900813 (2019).

61. Yousefian, H. et al. Beyond acid treatment of PEDOT:PSS: decoding mechanisms of electrical conductivity enhancement. Mater. Adv. 5, 4699–4714 (2024).

62. Bießmann, L. et al. Highly Conducting, Transparent PEDOT:PSS Polymer Electrodes from Post-Treatment with Weak and Strong Acids. Adv Elect Materials 5, 1800654 (2019).

63. Leaf, M. A. & Muthukumar, M. Electrostatic Effect on the Solution Structure and Dynamics of PEDOT:PSS. Macromolecules 49, 4286–4294 (2016).

64. Feig, V. R., Tran, H., Lee, M. & Bao, Z. Mechanically tunable conductive interpenetrating network hydrogels that mimic the elastic moduli of biological tissue. Nat Commun 9, 2740 (2018).

65. Yang, J. et al. 3D-Printed highly stretchable conducting polymer electrodes for flexible supercapacitors. J. Mater. Chem. A 9, 19649–19658 (2021).

66. Kan, X. et al. Granular hydrogels with tunable properties prepared from gum Arabic and protein microgels. International Journal of Biological Macromolecules 273, 132878 (2024).

67. Zhang, S. et al. Room-Temperature-Formed PEDOT:PSS Hydrogels Enable Injectable, Soft, and Healable Organic Bioelectronics. Advanced Materials 32, 1904752 (2020).

68. Wan, R. et al. 3D printing of highly conductive and strongly adhesive PEDOT:PSS hydrogel-based bioelectronic interface for accurate electromyography monitoring. Journal of Colloid and Interface Science 677, 198–207 (2025).

69. Wan, R. et al. A reusable, healable, and biocompatible PEDOT:PSS hydrogel-based electrical bioadhesive interface for high-resolution electromyography monitoring and time–frequency analysis. Chemical Engineering Journal 490, 151454 (2024).

70. Wakabayashi, T., Katsunuma, M., Kudo, K. & Okuzaki, H. pH-Tunable High-Performance PEDOT:PSS Aluminum Solid Electrolytic Capacitors. ACS Appl. Energy Mater. 1, 2157–2163 (2018).

71. Pérez, L. A., Hernández, R., Alonso, J. M., Pérez-González, R. & Sáez-Martínez, V. Granular Disulfide-Crosslinked Hyaluronic Hydrogels: A Systematic Study of Reaction Conditions on Thiol Substitution and Injectability Parameters. Polymers 15, 966 (2023).

72. Tyliszczak, B., Drabczyk, A., Kudłacik-Kramarczyk, S., Bialik-Wąs, K. & Sobczak-Kupiec, A. In vitro cytotoxicity of hydrogels based on chitosan and modified with gold nanoparticles. J Polym Res 24, 153 (2017).

73. Vilinsky, I. & Johnson, K. G. Electroretinograms in Drosophila: a robust and genetically accessible electrophysiological system for the undergraduate laboratory. J Undergrad Neurosci Educ 11, A149–157 (2012).

74. Nishino, H., Domae, M., Takanashi, T. & Okajima, T. Cricket tympanal organ revisited: morphology, development and possible functions of the adult-specific chitin core beneath the anterior tympanal membrane. Cell Tissue Res 377, 193–214 (2019).

75. Anderson, M. J. et al. The “Smellicopter,” a bio-hybrid odor localizing nano air vehicle. in 2019 IEEE/RSJ International Conference on Intelligent Robots and Systems (IROS) 6077–6082 (IEEE, Macau, China, 2019). doi:10.1109/IROS40897.2019.8968589.

76. Gupta, P. et al. Augmenting insect olfaction performance through nano-neuromodulation. Nat. Nanotechnol. 19, 677–687 (2024).

77. Lu, Y. & Liu, Q. Insect olfactory system inspired biosensors for odorant detection. Sens. Diagn. 1, 1126–1142 (2022).

78. Kim, B. et al. Olfactory receptor neurons generate multiple response motifs, increasing coding space dimensionality. eLife 12, e79152 (2023).

79. White, P. R. The electroantennogram response: Effects of varying sensillum numbers and recording electrode position in a clubbed antenna. Journal of Insect Physiology 37, 145–152 (1991).

80. Zhao, J., Li, Z., Zhao, Z., Yang, Y. & Yan, S. Electroantennogram reveals a strong correlation between the passion of honeybee and the properties of the volatile. Brain Behav 10, e01603 (2020).

81. Baker, T. C. & Roelofs, W. L. Electroantennogram responses of the male moth, Argyrotaenia velutinana to mixtures of sex pheromone components of the female. Journal of Insect Physiology 22, 1357–1364 (1976).

82. Zhou, T. et al. 3D printable high-performance conducting polymer hydrogel for all-hydrogel bioelectronic interfaces. Nat. Mater. 22, 895–902 (2023).

83. Ghasemi-Mobarakeh, L. et al. Application of conductive polymers, scaffolds and electrical stimulation for nerve tissue engineering. J Tissue Eng Regen Med 5, e17–e35 (2011).

84. Shi, M., Dong, R., Hu, J. & Guo, B. Conductive self-healing biodegradable hydrogel based on hyaluronic acid-grafted-polyaniline as cell recruitment niches and cell delivery carrier for myogenic differentiation and skeletal muscle regeneration. Chemical Engineering Journal 457, 141110 (2023).

85. International Organization for Standardization. Biological Evaluation of Medical Devices Part 5: Tests for in Vitro Cytotoxicity. https://www.iso.org/standard/36406.html (2009).

