## Supplementary Information for "PEDOT:PSS Microparticles for Extrudable and Bioencapsulating Conducting Granular Hydrogel Bioelectronics"

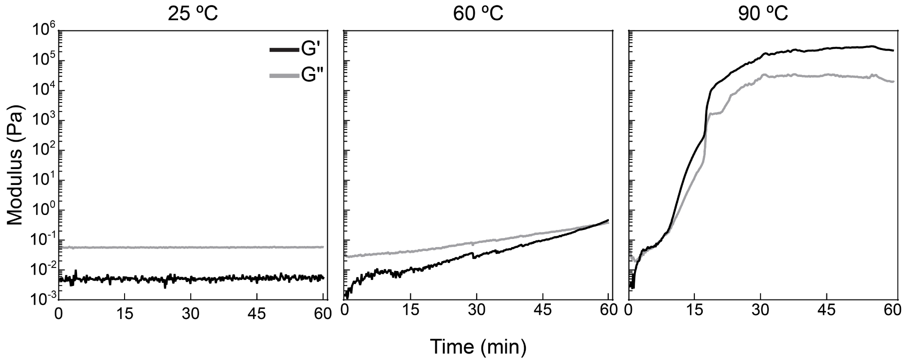


**Supplementary Figure 1. Repeated runs of oscillatory shear rheology of PEDOT:PSS colloidal aqueous dispersion at designated temperatures.** No gelation (G'-G" cross-over) was observed at 25 ºC, nor changes in G' and G". Gelation occurred slowly at 60 ºC with the gel point at 57.64 minutes. The liquid-to-gel transition (G'-G" cross-over) occurred at 90 ºC at 2.49 minutes (1% strain, 1 rad $s^{-1}$). Trends in data are similar to those shown in Figure 2.

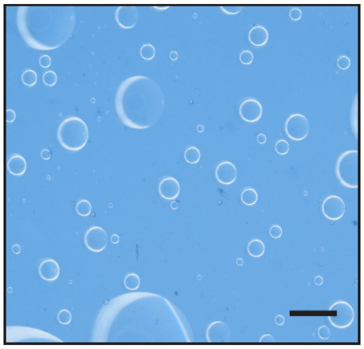

**Supplementary Figure 2. PEDOT:PSS droplets diffused into surrounding aqueous phase when isolation from oil phase was attempted.** Water-in-oil emulsions performed at room temperature did not result in aqueous stable microparticles due to unsuccessful crosslinking as evidenced by lack of observable PEDOT:PSS droplets upon introduction of water during washing process. Only mineral oil droplets were observed in deionized water. Scale bar 100 μm.

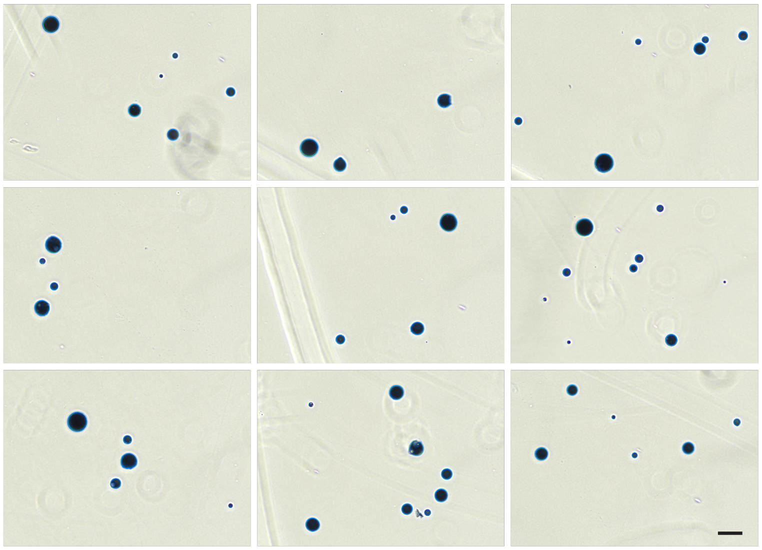

**Supplementary Figure 3. Images of microparticles fabricated using 90 ºC oil phase.** Additional images of microparticles fabricated using 90 ºC oil phase demonstrate the uniformity of the microparticle shape, translucence, and color. Scale bar 100 µm.

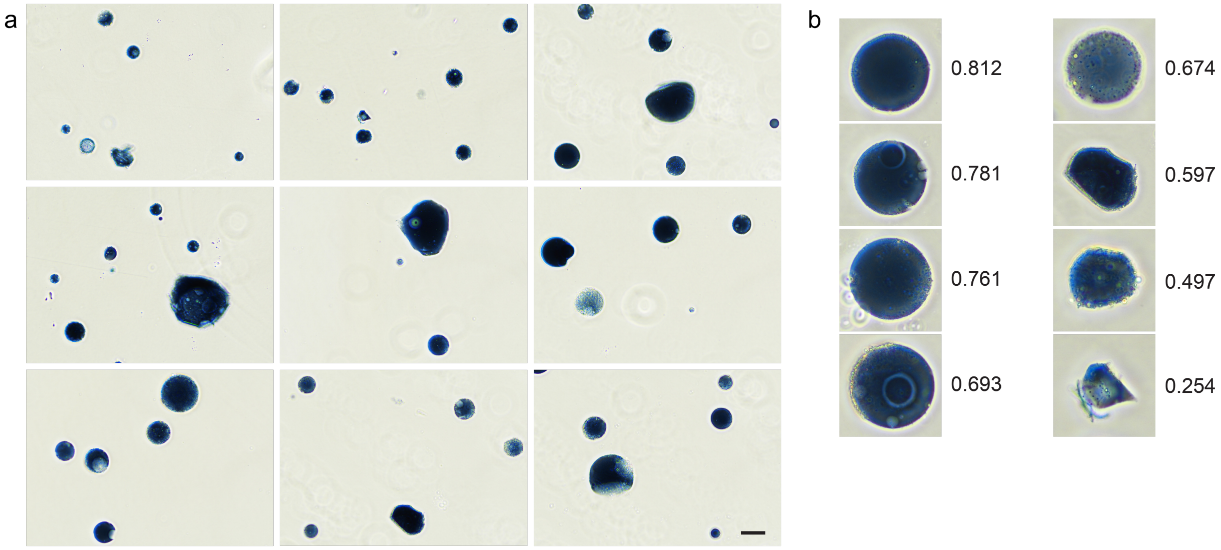


**Supplementary Figure 4. Images of microparticles fabricated using 60 ºC oil phase.** A) Additional images of microparticles fabricated using 60 ºC oil phase demonstrate some irregular shapes and translucence of some microparticles. Scale bar 100 µm. B) While spherical and smooth bordered particles are observed (left), irregular shapes and particle borders (right) are also observed and contribute substantially to the low mean circularity of the microparticle population. Microparticles shown in images from 4a repeated in 4b to demonstrate range of circularities measured in the population.

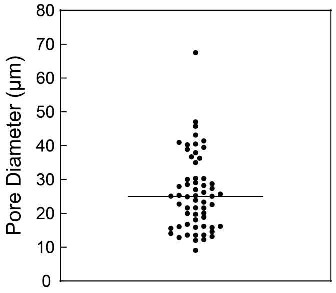


**Supplementary Figure 5. PEDOT:PSS granular hydrogel possesses pores of similar sizes to cells.** PEDOT:PSS granular hydrogel made from densely packed microparticles possessed pores primarily between 10 µm and 50 µm. N=30 pores taken from 2 granular hydrogel samples, 15 pores measured at random per sample. Mean pore diameter represented with line.

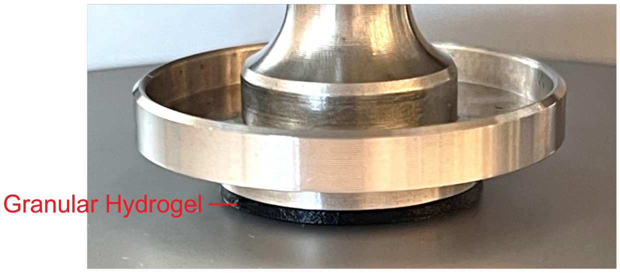


**Supplementary Figure 6. Loading of PEDOT:PSS granular hydrogel on rheometer.** The densely packed PEDOT:PSS microparticles were loaded onto the rheometer Peltier plate, and the 20 mm parallel plate was lowered onto the material at a gap height of 1100 µm. This resulted in the formation of an even disc of material.

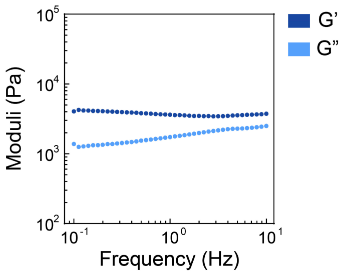


**Supplementary Figure 7. Frequency sweep of PEDOT:PSS granular hydrogel.** A frequency sweep was conducted at 1% strain. The storage modulus of the material exhibited linear behavior between 0.1 and 10 Hz. A frequency of 1 Hz was used in all further testing.

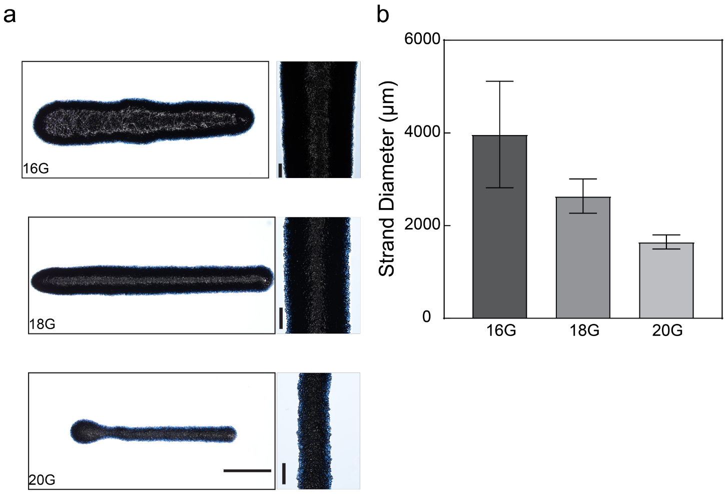


**Supplementary Figure 8. PEDOT:PSS granular hydrogel can be extruded via 3D printing and maintains shape of 3D printed strands.** A) Extrusion of the PEDOT:PSS granular hydrogel was achieved through 16, 18, and 20 G tapered tips. Scale bars 5000 µm for whole strand view and 1000 µm for insets. B) The width of 3D printed strands was quantified at nine equally spaced points across the strands and an average width for each strand was calculated. A 58.47% reduction in width was achieved by using a 20 G tapered tip in comparison to the 16 G tapered tip. N=3 strands per tapered tip. Mean and standard deviation presented.

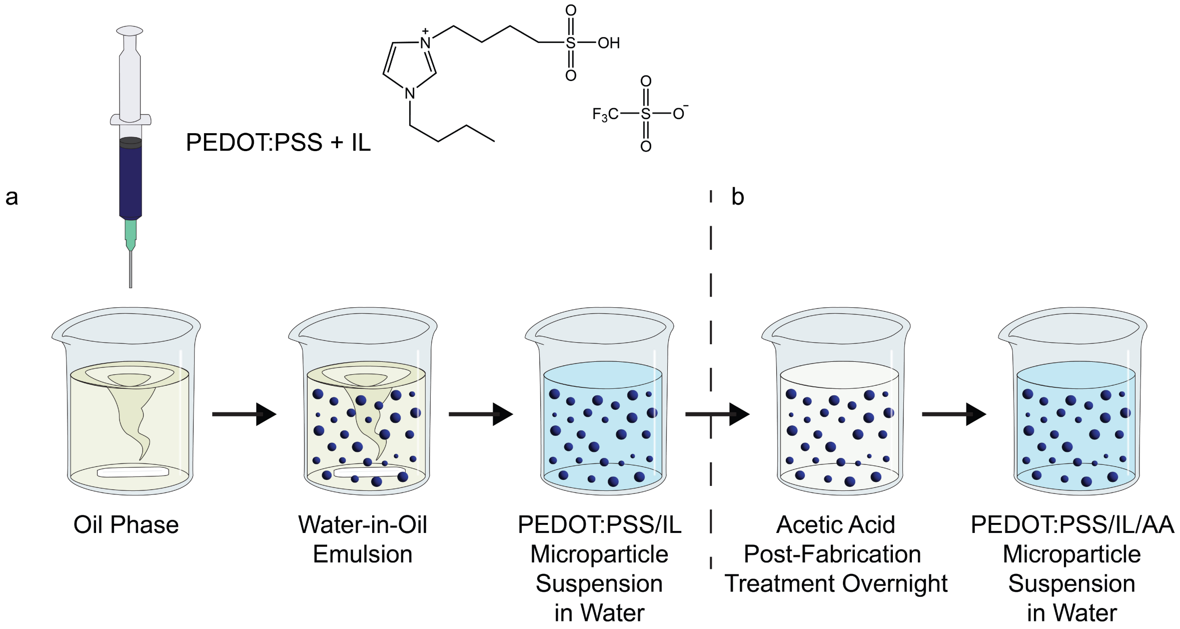


**Supplementary Figure 9. Methods of making PEDOT:PSS microparticles with inclusion of ionic liquid and post-fabrication treatment with acetic acid.** A) Ionic liquid (IL) was incorporated into the PEDOT:PSS colloidal aqueous dispersion which was then added to the 90 ºC oil phase being stirred at 750 RPM. The emulsion was maintained for 60 minutes. B) After isolation from the oil phase and water washing, PEDOT:PSS microparticles were then suspended in 17.3 M acetic acid overnight. Additional water washing was completed following post-treatment to remove acetic acid.

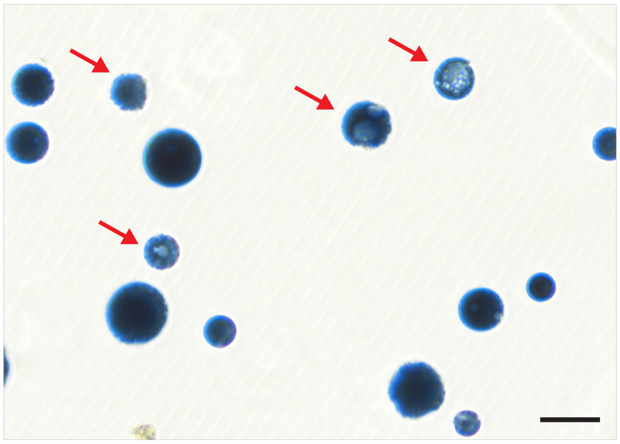

**Supplementary Figure 10. PEDOT:PSS microparticles prepared solely from PEDOT:PSS colloidal aqueous dispersion (no ionic liquid) and with acetic acid post-treatment have heterogeneous appearance throughout the population.** PEDOT:PSS/AA microparticles displayed changes in microparticle translucency/color and rougher particle borders which could indicate material loss. Example microparticles that showcased such changes are indicated with red arrows. Scale bar 100 μm.

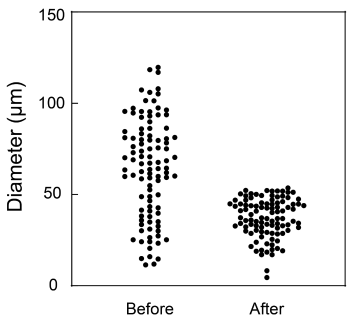


**Supplementary Figure 11. Filtration of microparticle suspensions can be used to define microparticle diameter range of population.** Filtration of PEDOT:PSS/IL/AA microparticles through a cell strainer with 60 μm pores reduced the maximum microparticle diameter from 120 μm to 54 μm.

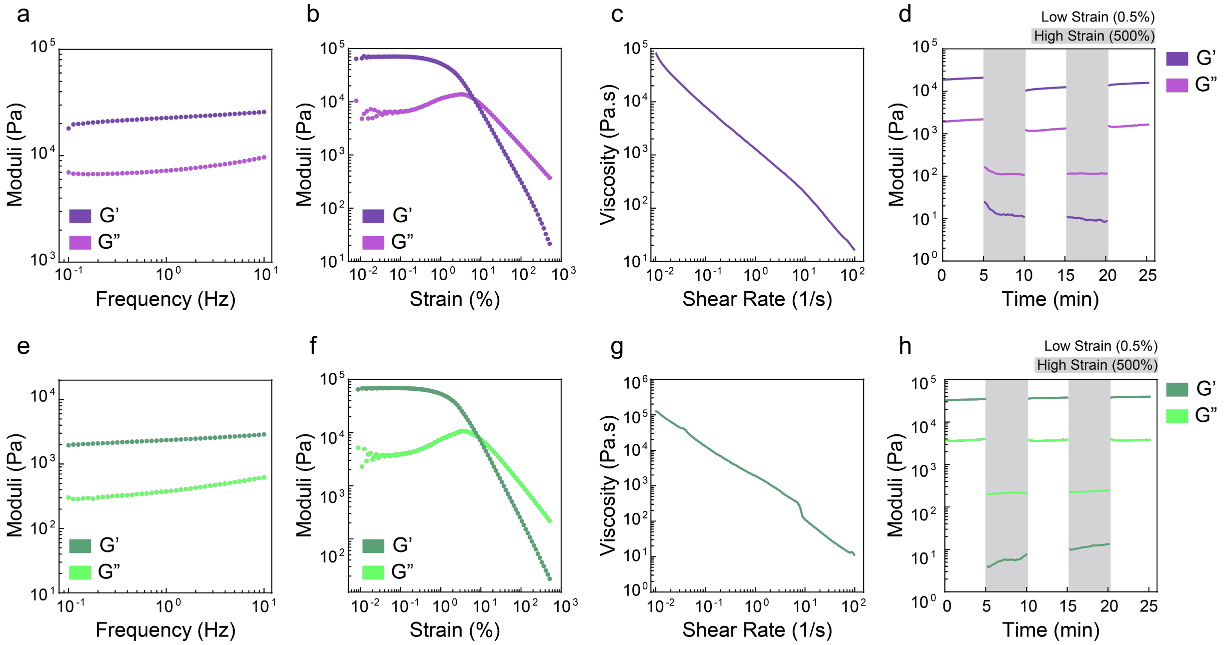


**Supplementary Figure 12. PEDOT:PSS/IL and PEDOT:PSS/IL/AA granular hydrogels exhibit shear-thinning and self-healing dynamic mechanical properties.** Oscillatory shear and rotational rheology of PEDOT:PSS/IL (top, purple) and PEDOT:PSS/IL/AA (bottom, green) granular hydrogels. A,E) The storage moduli of the densely packed PEDOT:PSS/IL and PEDOT:PSS/IL/AA microparticles exhibited linear behavior between 0.1 and 10 Hz (1% strain). A frequency of 1 Hz was used in all further testing. B,F) The densely packed microparticle populations exhibited deformation and underwent gel-to-liquid transition as evidenced by decreasing moduli and a G'-G" crossover (1 Hz). C,G) Shear-thinning behavior of the densely packed microparticle populations was confirmed as evident by a decrease in viscosity in response to increasing shear rate. D,H) A time sweep with alternating applications of low and high strain indicated recovery of the material (G'>G") at low strains following high strain (1 Hz).

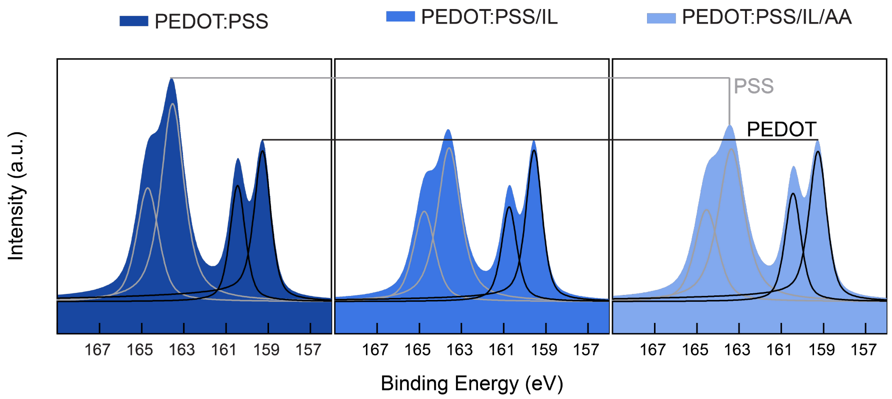


**Supplementary Figure 13. X-ray photoelectron spectroscopy (XPS) of PEDOT:PSS microparticles indicates decreasing PSS content with the inclusion of IL and further post-treatment with acetic acid.** PEDOT:PSS, PEDOT:PSS/IL, and PEDOT:PSS/IL/AA microparticles were evaluated via XPS for PSS content relative to that of PEDOT. The de-convoluted profiles were fitted with asymmetric functions and the PSS/PEDOT ratio was calculated from a comparison of the area under the curves of the XPS fits (representative plots of data presented in Figure 6).

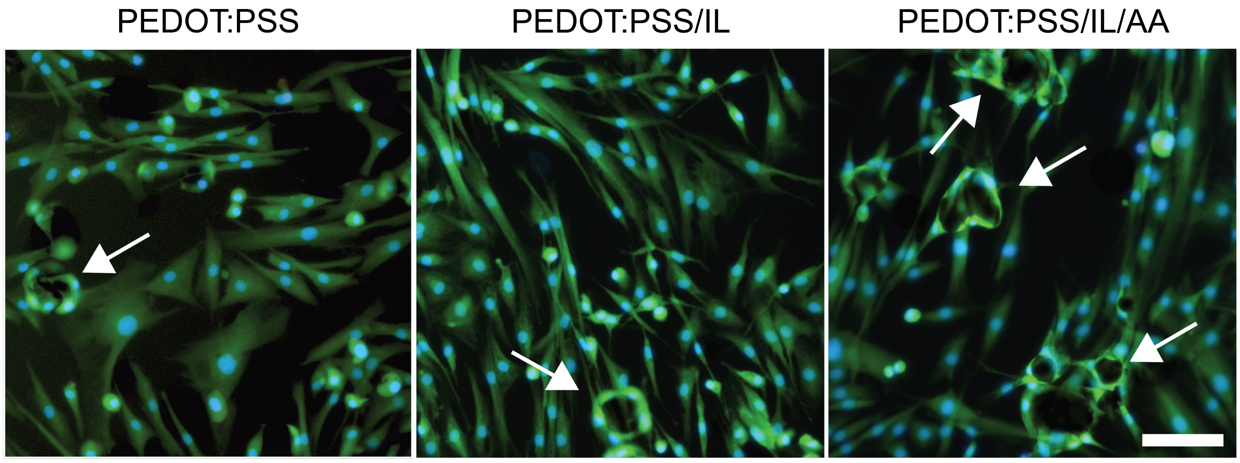


**Supplementary Figure 14. Fibroblasts wrap around microparticles.** Fibroblasts were observed to have wrapped around individual microparticles (arrows) after only 24 hours of contact between the microparticles and cell monolayers. Scale bar 100 µm.

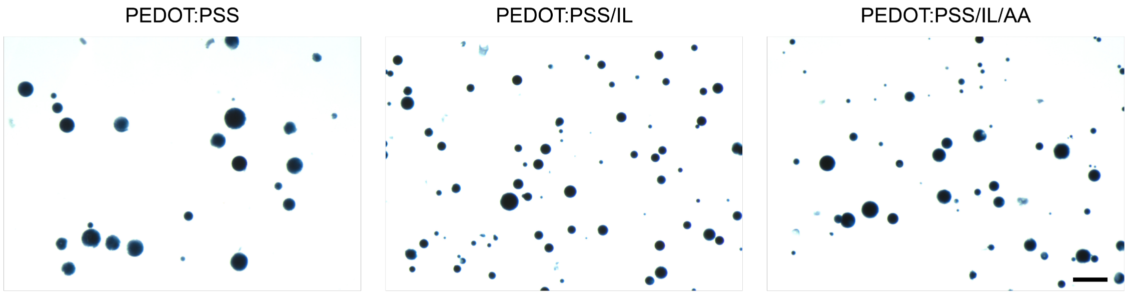


**Supplementary Figure 15. PEDOT:PSS, PEDOT:PSS/IL, and PEDOT:PSS/IL/AA microparticles exhibit long-term aqueous stability.** Microparticles in deionized water for at least 30 days show no visible changes indicative of material degradation. Scale bar 200 µm.

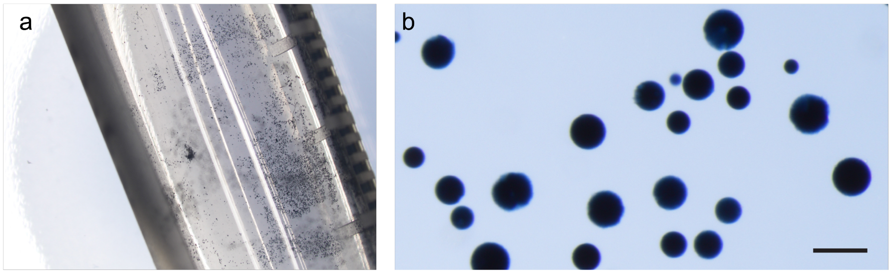


**Supplementary Figure 16. Microparticles demonstrate potential for long-term storage.** Microparticles were frozen at -20 ºC and lyophilized. A) Microparticles did not form aggregates during lyophilization in a micro-tube. B) Microparticles were rehydrated in water and no obvious differences were observed. Scale bar 100 µm.
